# An oligodendrocyte silencer element underlies the pathogenic impact of lamin B1 structural variants

**DOI:** 10.1101/2023.08.03.551473

**Authors:** Bruce Nmezi, Guillermo Rodriguez Bey, Talia DeFrancesco Oranburg, Kseniia Dudnyk, Santana M. Lardo, Nathan Herdman, Anastasia Jacko, Sandy Rubio, Emanuel Loeza Alcocer, Julia Kofler, Dongkyeong Kim, Julia Rankin, Emma Kivuva, Nicholas Gutowski, Katherine Schon, Jelle van den Ameele, Patrick F. Chinnery, Sérgio B. Sousa, Filipe Palavra, Camilo Toro, Filippo Pinto e Vairo, Jonas Saute, Lisa Pan, Murad Alturkustani, Robert Hammond, Francois Gros-Louis, Michael Gold, Yungki Park, Geneviève Bernard, Raili Raininko, Jian Zhou, Sarah J. Hainer, Quasar S. Padiath

## Abstract

The role of non-coding regulatory elements and how they might contribute to tissue type specificity of disease phenotypes is poorly understood. Autosomal Dominant Leukodystrophy (ADLD) is a fatal, adult-onset, neurological disorder that is characterized by extensive CNS demyelination. Most cases of ADLD are caused by tandem genomic duplications involving the lamin B1 gene (*LMNB1*) while a small subset are caused by genomic deletions upstream of the gene. Utilizing data from recently identified families that carry *LMNB1* gene duplications but do not exhibit demyelination, ADLD patient tissues, CRISPR modified cell lines and mouse models, we have identified a novel silencer element that is lost in ADLD patients and that specifically targets overexpression to oligodendrocytes. This element consists of CTCF binding sites that mediate three-dimensional chromatin looping involving the *LMNB1* and the recruitment of the PRC2 repressor complex. Loss of the silencer element in ADLD identifies a previously unknown role for silencer elements in tissue specificity and disease causation.

## Introduction

Many neurological disorders are caused by mutations in genes that are expressed in multiple cell types. Why the CNS is preferentially impacted in these diseases has been a longstanding puzzle. The fatal, adult onset, progressive neurological disorder Autosomal Dominant Leukodystrophy (ADLD, OMIM# 169500) is one such disease. Symptoms begin around the fifth or sixth decade of life with the primary pathology being widespread CNS demyelination with patients rarely living beyond their mid 70s^1–4^.

Most cases of ADLD are caused by tandem genomic duplications (ADLD-Dup) involving the lamin B1 gene (*LMNB1*) while a smaller subset are caused by genomic deletions (ADLD-Del) upstream of the gene^1,4–7^. In both cases, the pathological variants are completely penetrant and increased expression of LMNB1 is thought to underlie the disease phenotype^3,6,7^. Lamin B1 is an intermediate filament protein that is an integral part of the nuclear lamina, a fibrous meshwork found adjacent to the inner nuclear membrane in most mammalian cells^8,9^. It is critical for the proper maintenance of nuclear architecture and multiple other cellular processes^8,9^. Lamin B1 is widely expressed in many different cell types and tissues^10,11^ and why duplications or upstream deletions involving this gene causes a specific CNS demyelinating phenotype is a mystery ^4^. In addition, whether duplications and upstream deletions share a common mechanism to cause lamin B1 overexpression is unknown^4^.

Also unclear are the specific cell types that are involved in disease process. We have previously demonstrated that oligodendrocyte (OLs), the cell type that produces myelin in the CNS, are critical to the disease process using a transgenic mouse where we targeted human FLAG-tagged LMNB1 overexpression to OLs using a *Plp1* promoter construct^12^, a gene that is highly expressed in mature oligodendrocytes^13,14^. These *PLP-LMNB1* transgenic (TG) mice exhibited severe vacuolar degeneration of the white matter resulting in late age onset motor dysfunction, muscle wasting, paralysis and did not survive beyond 13 months^13^. The vacuolar degeneration was very similar to the pathological changes observed in human ADLD brain sections while the age-dependent motor dysfunction was reminiscent of symptoms in ADLD patients^15,16^. Whether or how other cell types such as astrocytes, microglia or neurons contribute to the demyelination phenotype is still unclear.

In this report, we present the identification of three independent families with multiple members segregating tandem duplications involving the *LMNB1* gene, but strikingly, that do not demonstrate any demyelination phenotype, even at an advanced age. These data suggest that merely having an extra copy of *LMNB1* does not result in ADLD and implicates other mechanisms regulating the overexpression. Using a combination of data derived from disease and non-disease causing *LMNB1* structural variants, ADLD patient tissues, CRISPR editing of diverse cell lines and mouse models, we have identified a novel silencer element that regulates the expression of lamin B1 specifically in oligodendrocytes. The loss of this silencer element in ADLD provides a parsimonious mechanism explaining tissue specificity and how both the duplication and upstream deletion can lead to lamin B1 overexpression and furthers our understanding of the role of non-coding regulatory elements in disease causation.

## Results

### Large tandem genomic duplications involving *LMNB1* do not cause ADLD

Genomic duplications of chr5q23.2, encompassing *LMNB1*, were initially identified in three independent multi-generational families (families F1-F3, Supplementary Fig. 1a-c) by whole genome wide array comparative genomic hybridization (aCGH) studies. None of the adult subjects studied exhibited symptoms of ADLD even though they were much older than the age of onset for the disease, the ages of adult subjects ranged from 47 to 82 years (Detailed clinical descriptions not presented, available upon request). Magnetic resonance imaging (MRI), a key tool for the diagnosis of leukodystrophies, revealed that none of these individuals exhibited leukodystrophic changes (Fig 1a-f, MRI data not shown for all individuals). MRI abnormalities have been shown to precede neurological symptoms in ADLD (Fig.1m-r) and can be identified as early as 29 years of age^17^. The fact that the older subjects with the duplications did not exhibit any MRI abnormalities or significant disability even in their mid 70s (Fig. 1a-f), an age at which no ADLD patients live beyond, further supports that they do not suffer from ADLD. Interestingly, although individuals did not exhibit the demyelination phenotype, some of them exhibited symptoms such as intermittent bowel or bladder dysfunction or orthostatic hypotension which can both be part of the early autonomic symptoms in ADLD.

**Figure 1:**
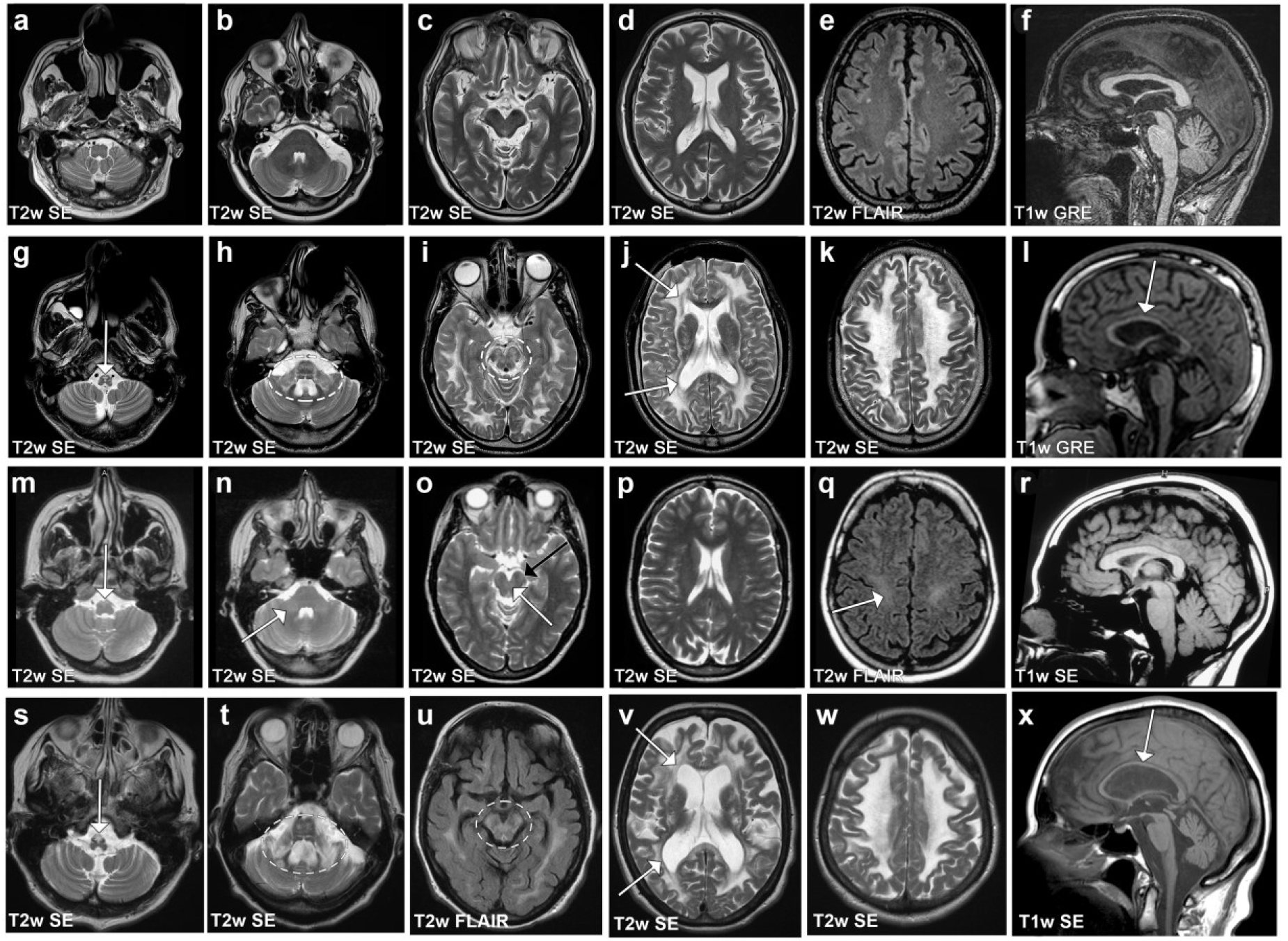
MR image series of subjects with *LMNB1* duplications. **a-f** Subject F2-1 at age 75 years exhibiting no leukodystrophic changes. The small cerebral white matter hyperintensities in **e** are nonspecific and may be age-related. **g-l** Subject F4-1 at age 34 years showing a pathologic high T2 signal intensity in the pyramids of the medulla oblongata (**g**, arrow), in the pons and middle cerebellar peduncles (**h**), mesencephalon (**i**), and in all cerebral lobes (**i-k**). A less affected periventricular rim in T2w SE images (**j**, arrows) is characteristic for *LMNB1*-related leukodystrophy. The corpus callosum is thin (**l**, arrow). **m-r** A subject with a canonical ADLD-causing *LMNB1* duplication at the age 35 when they were still asymptomatic but had a mild T2 signal intensity increase in the pyramids (**m**, arrow), in the middle and upper cerebellar peduncles (**n-o**, white arrows), and in the corticospinal tracts both in the mesencephalic **(o**, black arrow) and in the uppermost parts **(q**, arrow). **s-x** A subject with a canonical ADLD-causing *LMNB1* duplication and clinical disease at the age 65 exhibits similar MR abnormalities to F4-1. A pathologic high T2 signal intensity in the pyramids of the medulla oblongata (**s**, arrow), in the pons and middle cerebellar peduncles **(t**), mesencephalon (**u**), and in all cerebral lobes (**u-w**). A less affected periventricular rim is seen **(v**, arrows) and the corpus callosum is thin **(x**, arrow). Abbreviations: T2w SE = T2-weighted, T1w = T1-weighted, SE = spin echo sequence, FLAIR = fluid attenuated inversion recovery sequence, GRE = gradient echo sequence.

Subjects with the *LMNB1* duplications were subsequently analyzed with a custom high resolution CGH array that we had previously designed for the *LMNB1* region^13^ which allowed us to identify precise duplication junction boundaries (Supplementary Fig. 2, Fig. 2a, Supplementary table 1). Sequencing PCR products using primers that flanked the duplication junctions revealed that they were in a tandem head to tail configuration (Supplementary Fig. 3, Fig. 2a), similar to the ADLD-causing duplications^5^.

**Figure 2:**
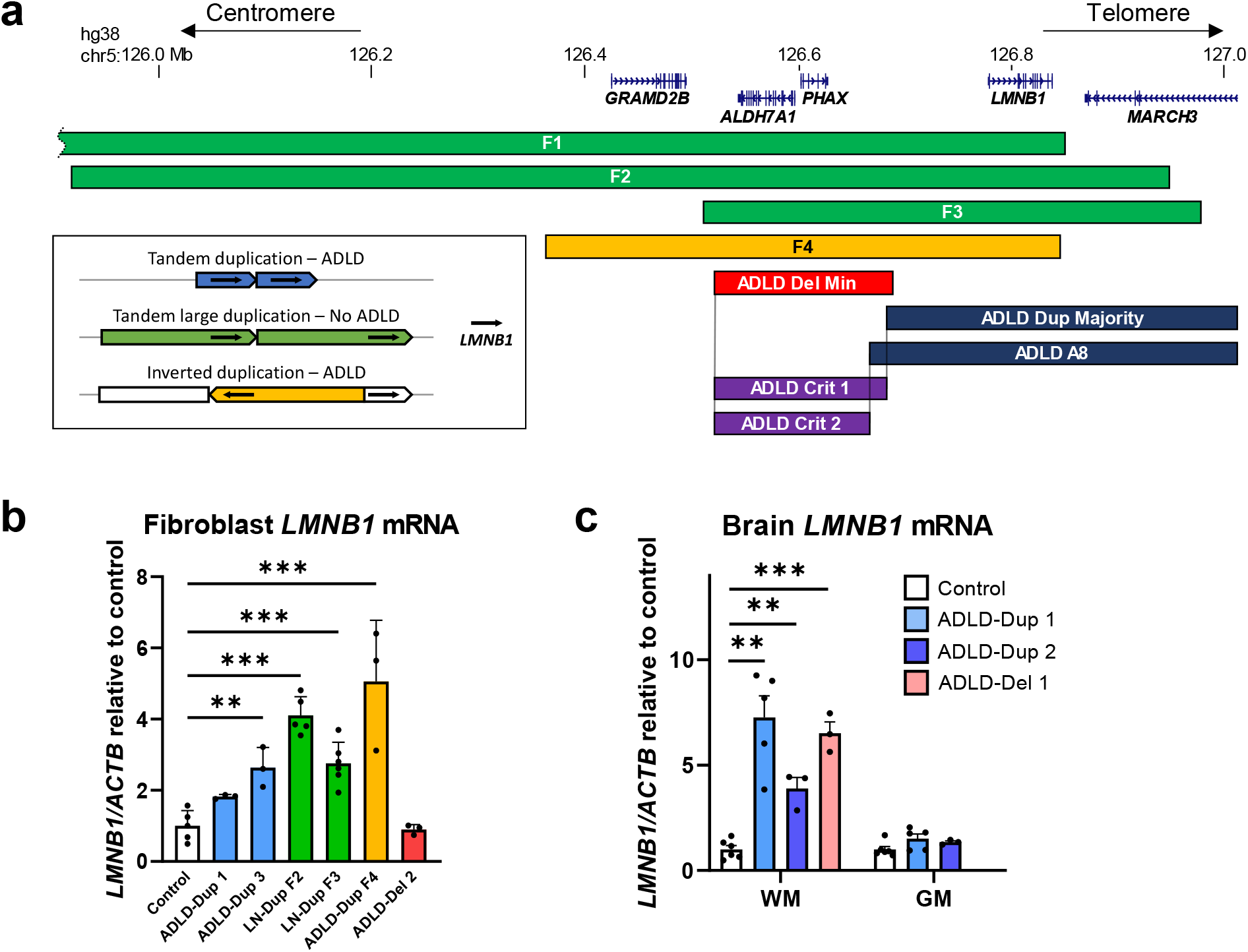
*LMNB1* structural variants and expression analysis. **a** Schematic of duplication extents in LN-Dup (F1-F3, green) and F4 (yellow) families. For the canonical disease-causing ADLD duplication (blue), duplication extents were composites based on previous reports^5^ where precise duplication boundaries were identified. ADLD-Dup majority (dark blue) represents the maximal extents of all disease-causing duplications from multiple families^5^. In one family, A8^5^ (dark blue), the centromeric end of the duplication extended beyond the ADLD-Dup majority and is shown separately. ADLD-Del Min (red) is a composite of multiple patients with deletions and represents the smallest deleted region causing ADLD^6,7^. ADLD-Crit 1 and ADLD-Crit 2 represent the smallest critical regions that are included in the ADLD-Del Min and LN-Dup but not in ADLD-Dup Majority or A8, respectively. Inset box shows the orientation of the various structural variants, with *LMNB1* depicted as a black arrow. Both ADLD Dup and LN-Dup (F1-F3) have a duplicated *LMNB1* gene with a similar head-to-tail tandem configuration, but LN-Dups are larger and contain more of the *LMNB1* upstream regulatory region, towards the centromeric end. The F4 duplication is inverted and inserted in the original duplicated segment as previously identified^5^. **b** Fibroblasts from both LN-Dup and ADLD-Dup patients show significantly increased *LMNB1* expression compared to controls, as measured by real time PCR relative to *GAPDH*. No difference is observed in *LMNB1* expression between ADLD-Del2 and controls. **c** ADLD-Dup and Del patients show significantly higher expression of *LMNB1* in white matter vs. grey matter in comparison to control brain samples. mRNA levels were measured by real time PCR relative to *GAPDH*. Note that we did not have grey matter sample for the ADLD-Del-1 patient. ** - p < 0.01. *** - p < 0.001. All comparisons are between controls and LN-Dup or ADLD samples using t-tests. n ≥ 3 biological replicates for control brain and fibroblast samples. For ADLD-dup, ADLD-del and LN-Dup samples, n ≥ 3 replicates per sample. ADLD-Dup 1-3 and ADLD-Del 1-2 have been described previously ^5,6^.

The duplications in the three families ranged from 4.37 Mb (F1) and 1.03 Mb (F2) to 466 kb (F3) and did not share any common junction sequences suggesting that they were non-recurrent and arose independent of each other (Supplementary Fig. 3, Supplementary Table 1). We have termed these duplications ‘large normal duplications (LN-Dups)’, as they were not associated with a leukodystrophy phenotype. While these LN-Dups were also tandem, they were larger and extended more centromeric (Fig. 2a) than the canonical ADLD-Dups which ranged from 128 kb to 324 kb^5^.

The exception to this range was the F4-1 patient that we previously identified^5^ (Supplementary Fig. 1d, denoted as BR-1 in that publication) with a duplication of ∼475 kb involving *LMNB1* but whose clinical features had not been previously described. This patient did not have a head to tail tandem duplication but rather an inverted duplication (ADLD-Inv-Dup) that was inserted centromeric to the *LMNB1* gene (Fig. 2a)^5^. F4-1 exhibited, both clinically and radiologically, a more severe and accelerated form of ADLD with MRI features at age 34 (Fig 1g-l) resembling ADLD patients in their 60s with advanced stages of the disease (Fig. 1s-x) and was wheelchair bound at age 38 (Detailed clinical descriptions not presented, available upon request). Furthermore, both his mother and grandmother were reported to have suffered from a similar neurological disorder and died in their mid 30s (Supplementary Fig. 1).

These findings demonstrate that simply possessing an extra copy of the *LMNB1* is insufficient to cause the ADLD disease phenotype and that additional factors such as size and orientation of the duplication need to be considered when predicting the pathogenicity of *LMNB1* duplications.

### *LMNB1* is differentially expressed in ADLD patient tissues

We analyzed *LMNB1* expression from ADLD patients from whom we had fibroblast and brain samples (Fig. 2b, c). Two brain samples were obtained from ADLD patients with duplications and one from an ADLD patient with a deletion. The duplications in sample 1 and 2 were 213 kb and 278 kb while the sample 3 was from a patient with a deletion that encompassed a 608 kb region upstream of the *LMNB1* gene^5,15^. All three patients had typical ADLD presentations and the duplications and deletion have been described previously^5,6^.

RNA expression analysis from ADLD-Dup fibroblasts demonstrated an increase of ∼1.5 to 2-fold *LMNB1* expression, consistent with these cells having 3 vs. 2 copies of the *LMNB1* gene and similar to our previous results from other ADLD fibroblasts^5^ (Fig. 2b). However, the fibroblast line from the ADLD deletion patient did not exhibit any increase in *LMNB1* expression (Fig. 2b). Interestingly, fibroblasts derived from LN-Dup individuals also exhibited significant *LMNB1* overexpression comparable to ADLD-dup patients (Fig. 2b). Fibroblasts derived from the F4-1 patient exhibited the highest level of *LMNB1* expression (Fig. 2b).

*LMNB1* expression was then analyzed from ADLD brain samples compared to age and sex matched controls (Fig. 2c). We isolated RNA from the frontal grey matter, which is not affected in the disease and frontal white matter, which is a site of demyelination in ADLD. Intriguingly, while *LMNB1* expression in the grey matter did not exhibit statistically significant increases, expression in the white matter was increased from ∼4 to 7-fold in ADLD patients’ samples as compared to controls (Fig. 2c). This increase in the white matter expression of *LMNB1* is much higher than what would be expected from just an extra copy of the gene. As these are autopsy brain tissue it is possible that this increase of *LMNB1* expression could be due to alterations in other cell types secondary to the demyelination phenotype. However, it may also suggest a distinct transcriptional regulatory mechanism that directs the over expression specifically to the white matter where OLs are one of the predominant cell types.

### Oligodendrocytes are not uniquely susceptible to the pernicious effects of *LMNB1* overexpression

As OLs clearly drive a significant portion of the pathology in our mouse model^13^, two possibilities may exist to explain their preferential involvement in ADLD patients and the specificity of the demyelination phenotype. The first possibility is that these cells types are uniquely susceptible to the effects of *LMNB1* overexpression i.e. all cell types are exposed to similar *LMNB1* overexpression but only OLs suffer deleterious consequences. The second possibility is that the genomic rearrangements (ADLD-Dup and ADLD-Del) that underlie the disease result in a mis-expression of *LMNB1* that specifically targets OLs (and potentially other glial cells in the white matter) for maximal overexpression.

In addition to OLs in the CNS, the *Plp*1 promoter used in our TG mice also drives expression in Schwann cells^12^ that are responsible for peripheral myelination. We confirmed that the FLAG-tagged human *LMNB1* (*hLMNB1*) was also expressed in the sciatic nerve, part of the peripheral nervous system (Fig. 3a-c). If OLs are uniquely susceptible to the deleterious effects of *LMNB1*, we would expect peripheral myelination to be unaffected in our TG mice despite the overexpression of *LMNB1* in Schwann cells. We therefore studied the impact of *LMNB1* overexpression on myelination in sciatic nerves and discovered that the TG mice exhibited significant age dependent demyelination, as evidenced by reduction of Luxol Fast Blue (LFB) staining, a myelin specific stain (Fig.3d,e), loss of myelinated axons analyzed by electron microscopy analysis (Fig. 3f-h), and nerve conduction defects (Fig. 3i,j), when compared to wild type mice. These results indicate that overexpression of *LMNB1* in Schwann cells can also have deleterious effects and that OLs are not unique in their susceptibility to increased *LMNB1* levels. However, in ADLD patients, no peripheral nerve conduction defects have been described^1,17^ suggesting that peripheral myelination is unaffected. Combining the data from the previous two sections suggests that in ADLD patients, specific OL involvement is more likely due to targeted overexpression to the CNS.

**Figure 3:**
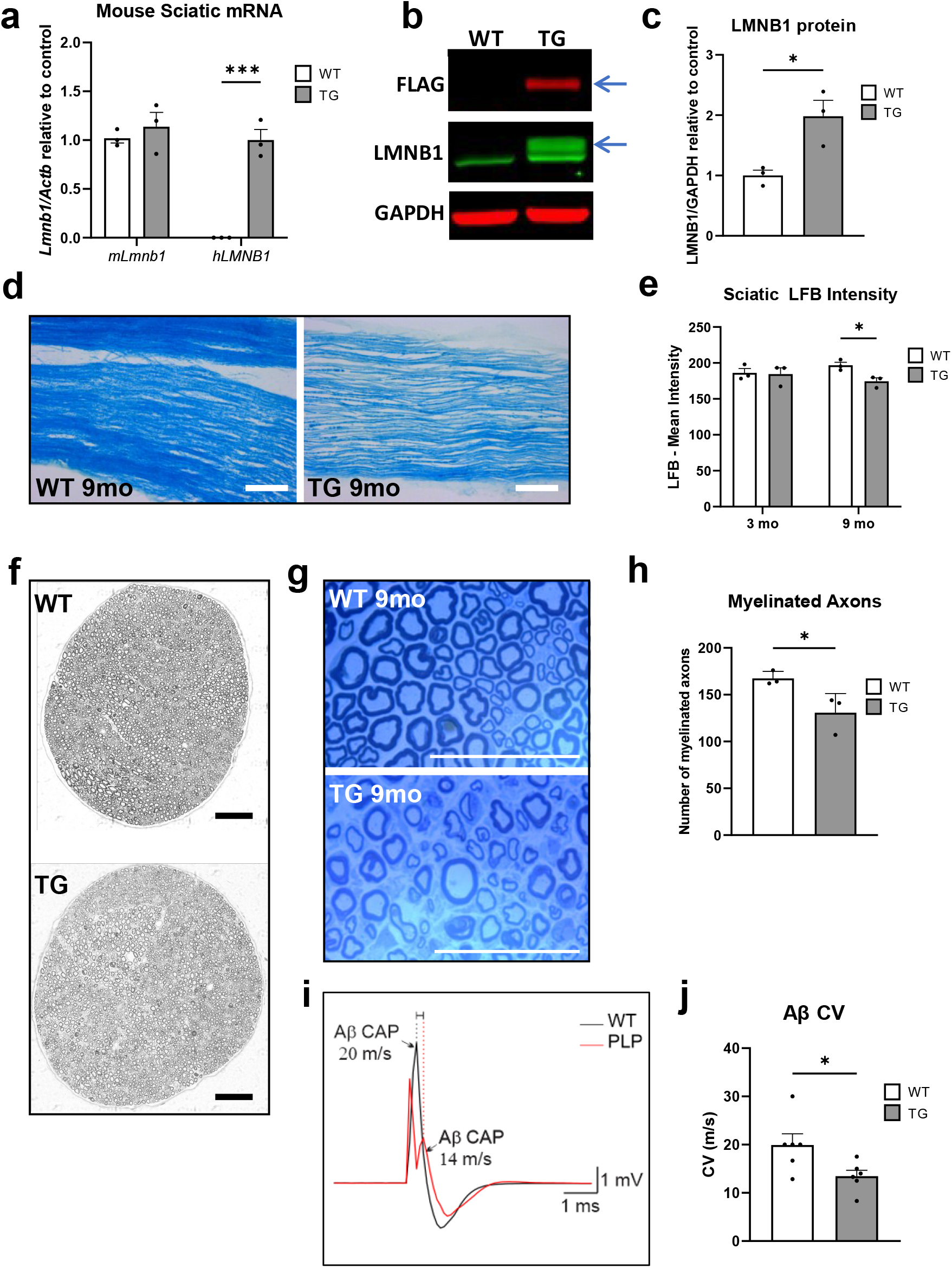
Peripheral nerve degeneration in LMNB1 overexpressing TG mice. **a** Real time PCR analysis from sciatic nerve samples from TG and WT mice demonstrate expression of the exogenous human *LMNB1 (hLMNB1)* only in TG samples. Mouse *Lmnb1 (mLmnb1)* expression is not altered between WT and TG samples. n=3 independent mouse samples for each group, *** p <0.001 by t-test. **b** Western blot of sciatic nerves from WT and TG mice probed for endogenous mouse LMNB1, Flag-tagged exogenous hLMNB1, and GAPDH as a loading control. **c** Quantification of immunoblot demonstrating that total LMNB1protein (mLMNB1 and hLMNB1) is overexpressed in sciatic nerves from TG mice. n=3 independent mouse samples for each group, * p <0.05 using t-tests. **d** Representative brightfield images of 20μm thick longitudinal sections of sciatic nerves from 9-month-old mice stained with luxol fast blue (LFB), scale bars = 100μm. **e** Quantification of the LFB staining shows less LFB staining in nerves from 9-month TG mice when compared to WT controls. n=3 independent mouse samples for each group * p <0.05 by one-way ANOVA. **f** Representative TEM montages of semi-thin transverse sections of sciatic nerves from 9-month-old WT and TG mice stained with toluidine blue. **g** Representative zoomed in images of the sections in **f**, scale bars = 50μm. **h** Quantification of the number of myelinated axons. Sciatic nerves from 9-month-old TG mice have fewer myelinated axons than those from WT. n=3 independent mouse samples for each group, * p <0.05 using t-tests. **i** Representative traces of recorded compound action potential (CAP) of Aβ large diameter fibers from sciatic nerves from 9-month-old WT (black line) and TG (red line) mice. **j** Quantification of conduction velocity in sciatic nerves from 9-month-old mice demonstrates slower conduction velocity in Aβ fibers from TG mice. n ≥ 3 independent mouse samples for each group, * p <0.05 using t-tests. All error bars represent error of the mean of biological replicates.

### Analysis of *LMNB1* structural variants identifies the presence of a silencer element that down-regulates *LMNB1* expression specifically in OL lineage cells

How does one explain the absence of the leukodystrophy phenotype in LN-Dups while the ADLD-dup, ADLD-del and ADLD-Inv-Dup results in demyelination? In light of the finding that *LMNB1* overexpression in ADLD may be targeted to specific CNS tissues or cell types, one possible mechanism is that *LMNB1* is under the control of a silencer genomic element that maintains low levels of expression in OLs (Fig. 4a). Gene expression analysis studies of mouse OL lineage cells have revealed that *Lmnb1* expression is dramatically reduced during the maturation of oligodendrocyte precursor cells (OPCs) into mature OLs^18^ (Supplementary Fig. 4a, b).

**Figure 4:**
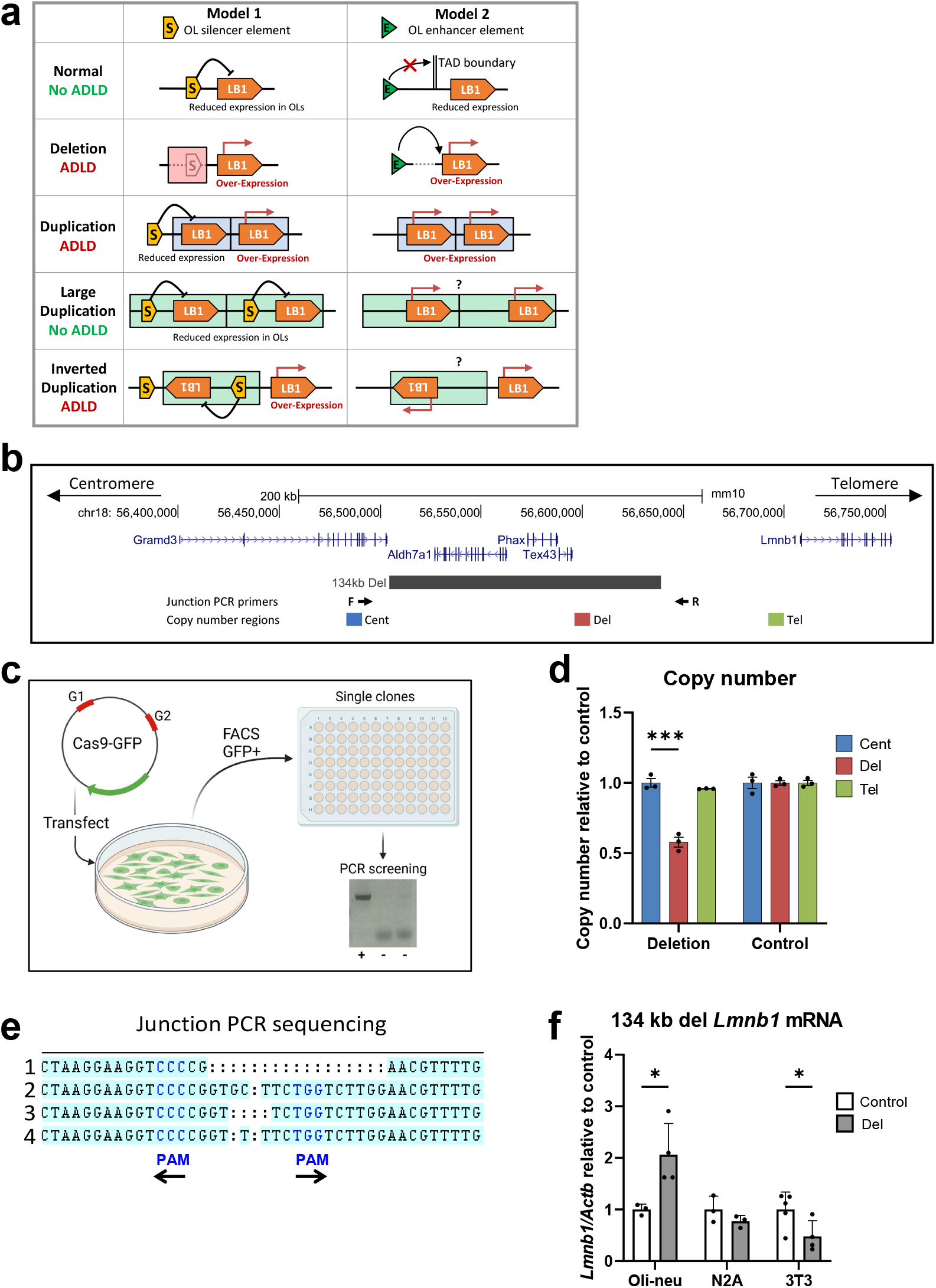
Silencer model for *LMNB1* overexpression in ADLD and generation of CRISPR/Cas9 mediated genomic deletions. **a** Model 1 - A silencer element acts to maintain low *LMNB1* expression in oligodendrocytes (OLs). In ADLD-Del, this silencer is deleted leading to overexpression. In ADLD-Dup, the *LMNB1* gene is duplicated but the silencer is not, also leading to overexpression in OLs. In LN-Dup, both the silencer element and *LMNB1* are duplicated resulting in the maintenance of reduced levels of *LMNB1* of the duplicated copy. In the inverted duplication, the inverted copy is inserted between the putative silencer element and its cognate *LMNB1* copy thereby preventing interaction with the silencer element. Model 2 – Based on a previous study of an ADLD-Del patient, proposes that the upstream deletion causes loss of a TAD boundary bringing an enhancer closer to *LMNB1* leading to overexpression^7^. However, it is unclear how this would explain why there is no disease due to LN-Dup and assumes an extra copy of the *LMNB1* gene is sufficient to produce disease. **b** UCSC genome browser view showing the syntenic ADLD-Crit 1 region in the mouse genome (134kb-Del, grey bar) deleted by CRISPR/Cas9. **c** Dual guide RNAs (G1 and G2) were cloned into a CRISPR/Cas9 plasmid followed by transfection and FACS sorting for GFP+ single cells. Deletion positive clones were identified by PCR screening using primers that amplify across the deletion junction. Only clones with deletions (+) will show a PCR product. **d** Copy number analysis of DNA from positive oli-neu clones with deletion using real time PCR demonstrates reduced copy number using primers within deleted region (red). n = 3 independent clones for, * p<0.05. Primers outside deleted region (blue and green) show copy number similar to control cells. **e** Sequencing deletion junctions using the junction PCR primers shown in **b** reveals each oli-neu clone has a unique sequence due to imperfect repair after CRISPR/Cas9 mediated deletions. Protospacer Adjacent Motif (PAM) sites are highlighted in blue. Note that PAM sites can be located on the reverse strand, with their orientation indicated by arrows. **f** *Lmnb1* mRNA expression as measured by real time PCR relative to bActin (*Actb*) is significantly higher in oli-neu cells with the deletion but is not significantly altered in N2A cells and reduced in 3T3 cells, compared to control cell lines. n ≥ 3 for both control and deletion clones for each cell type. *: p < 0.05, calculated using Student’s t test.

Based on our hypothesis, in the canonical ADLD-Dups, the *LMNB1* gene is duplicated while the silencer regulatory element is not. As a result, the duplicated copy of the *LMNB1* gene is no longer under the repressive control of the silencer element and overexpression occurs. In the case of the ADLD-Del, this silencer element is lost due to the upstream genomic deletion, leading to a similar overexpression (Fig. 4a, left panel). This mechanism would explain why the LN-Dups are not associated with demyelination as, in this case, the duplications also include the putative silencer element. The duplicated *LMNB1* gene is thus still repressed and would not exhibit a specific overexpression in OLs.

This model also explains the paradoxical case of the F4 duplication that results in ADLD, even though this duplication has a comparable size to the LN-Dups. Although it includes the putative silencer element, it is not arranged in a tandem orientation. Rather, the duplication is inverted and inserted between the silencer element and the non-duplicated copy of the *LMNB1* gene and could potentially result in the disruption of the spatial relationship between them (Fig. 4a). Why the F4 duplication results is a more severe phenotype can also be explained by our silencer model and is discussed in detail below.

An alternative model (Fig. 4a, right panel), that was proposed when the first ADLD-del family was identified, posited that the deletion resulted in the adoption of an alternative enhancer that acted on *LMNB1*^7^. However, this model assumed that the extra copy of *LMNB1* was sufficient to cause disease, which we now know is not the case. It cannot explain the lack of a disease phenotype in the LN-Dup families and why the F4 duplication results in a more severe form of ADLD.

Using the silencer model to explain the differential pathogenic effect of the *LMNB1* structural variants (SV), and by comparing the maximal centromeric extents of the ADLD-causing duplications and telomeric extents of ADLD-causing deletions to that of the centromeric extents of the nonpathogenic *LMNB1* duplications we were able to delineate a minimal critical genomic region that would be predicted to include this putative *LMNB1* suppressor element (Fig. 2a). We only utilized *LMNB1* ADLD-causing duplications whose exact duplication junctions were identified^5^. We noted that with the exception of one case, the centromeric extent of the majority of the ADLD duplications extended to genomic coordinates (hg38) chr5:126683196. The duplication in one family, A8, extended to genomic coordinates chr5:126667592^5^. These two boundaries allowed us to identify the ADLD critical regions 1 and 2, with sizes of ∼145 kb (ADLD Crit-1) and ∼160 kb (ADLD Crit-2), respectively (Fig. 2a). An analysis of this region in the mouse and human genome revealed a high degree of synteny with an almost complete conservation of gene order, suggesting that intergenic elements such as the putative silencer might also be conserved (Supplementary Fig. 5a, b).

To test whether the ADLD critical region influences lamin B1 expression, we carried out CRISPR/Cas9 mediated genomic deletions of a syntenic genomic segment in the mouse genome. This region in the mouse genome was identified using the ‘lift over’ function in the UCSC genome browser and corresponded to ∼134 kb (mm10, chr18: 56504521-56638837) on mouse chromosome 18 (Fig. 4b). Genomic deletions were generated in oli-neu, neuro2A (N2A) and NIH-3T3 (3T3) mouse cell lines that represented oligodendroglial, neuronal and fibroblast lineages, respectively. Undifferentiated oli-neu cells are considered similar to OPCs while differentiated oli-neu cells are thought to recapitulate OLs in early stages of maturity^19^. They have been extensively utilized in multiple studies as surrogates for early stage OLs, especially in analyzing epigenetic modifications and regulatory pathways in the OL lineage^20–26^. Relevant to our work, oli-neu cells also exhibit a significant reduction in *Lmnb1* expression levels upon differentiation similar to OLs, suggesting that the regulatory mechanisms that control *Lmnb1* expression are similar between primary OLs and oli-neu cells (Supplementary Fig. 4b).

At least three independent clones with a genomic deletion of ∼134 kb corresponding to the syntenic ADLD critical region were generated for each of the cell types. In all cases, clones were grown from single cells and presence of the deletions were confirmed by real time PCR to measure DNA copy number and sequencing of junction PCR products across the deletion breakpoints (Fig. 4c-e, Supplementary Fig. 6a, b). Sequencing deletion junctions also confirmed that each clone was unique as non-homologous end joining (NHEJ) repair of the breakpoints result in sequence ‘scars’ for each clone (Fig. 4e, Supplementary Fig. 6b). Strikingly, we observed a significant increase in *Lmnb1* expression levels in differentiated oli-neu cells with the 134 kb deletion but not in N2A or 3T3 cells (Fig. 4f). These results suggest that the ADLD critical region does indeed contain a regulatory element that can potentially act as a silencer in an oligodendrocyte specific manner. However, as the 134 kb deletion results in the disruption of the centromeric Topologically Associated Domain (TAD) boundary containing lamin B1 (Fig. 4a, right panel), the overexpression could still result from the action of a novel enhancer, as has been suggested previously^7^.

### Examination of the Lamin B1 topologically associated domain identifies a candidate regulatory region

To confirm the silencer hypothesis and narrow down this putative element we analyzed the 3D chromatin organization using previously published chromatin conformation capture data from human embryonic stem cells (hESC)^27^ (Fig.5a). Given that long range chromatin interactions preferentially occur within TADs^28,29^, we reasoned that the putative silencer would reside within the TAD encompassing the *LMNB1* gene. Two genomic regions located ∼121 kb centromeric to the *LMNB1* transcription and that were ∼19 kb apart were identified to interact with the human *LMNB1* gene (Fig. 5a). We also confirmed that the 3D genome organization was similar in mouse and human and that this interacting region was found in both genomes (Supplementary Fig. 5b). This 19 kb region is completely included in ADLD Crit 1 and partially included in ADLD Crit2, towards their telomeric ends, and was thus an attractive candidate for the *LMNB1* regulatory element (Fig. 5a). In addition, analysis of previously published PLAC-Seq data^30^ from human brain derived OLs identified genomic regions that interacted with the *LMNB1* promoter corresponding to the 19 kb candidate region we identified (Fig. 5b). This interaction was also present in human OLs based on single cell Hi-C experiments (Fig. 5e)^31^. Analysis of previous ATAC-Seq data from human OLs revealed chromatin accessibility peaks in the 19kb region (Fig. 5c)^30^. However, these regions were not enriched in H3K27Ac marks that are usually associated with enhancer elements (Fig. 5d). This would suggest that while this 19 kb region serves as a *LMNB1* regulatory element, it is unlikely to be an enhancer.

**Figure 5:**
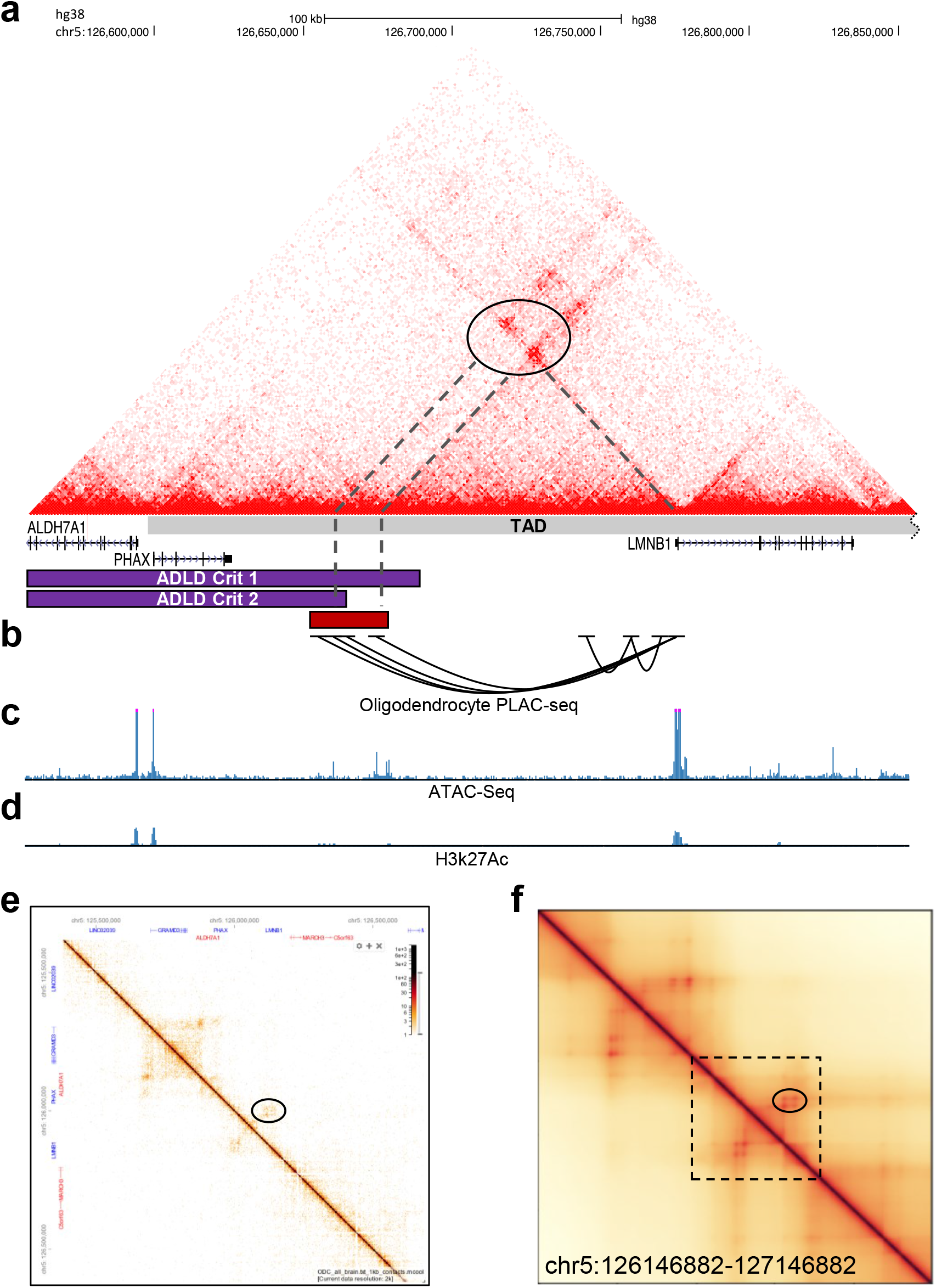
Analysis of 3D chromatin interactions identify a *LMNB1* regulatory element. **a** Micro-C chromatin interaction map^27^ (hg38) from human Embryonic Stem Cells (hESC) reveals distinct long-range interactions between the *LMNB1* promoter and upstream elements (dashed lines, oval). This 19 kb region (red bar) wholly overlaps ADLD Crit 1 and partially with ADLD Crit 2. The transcriptionally associated domain (TAD) containing *LMNB1* is also depicted. **b** PLAC-Seq maps from human OLs^30^ confirms the interaction between the *LMNB1* promoter and the 19 kb putative regulatory element. **c** ATAC seq data of human OLs^30^ identifies regions of open chromatin in the 19 kb region. **d** ChIP Seq data of human OLs^30^ reveals no enrichment of H3K27Ac in this region. **e** Single cell Hi-C data from human OLs^31^ revels a similar interaction between *LMNB1* and the 19 kb element (oval) as seen in **a**. **f** The interaction between *LMNB1* and the 19 kb regulatory element (oval) is recapitulated using the *Orca* simulation based on sequence data. Boxed region is the genomic segment depicted in **a**. All coordinates are from the hg38 human genome build.

### Interactions with the putative 19 kb regulatory element are perturbed by disease causing structural variants

To determine how the *LMNB1* SVs altered 3D genome organization and impacted the interaction between the 19 kb regulatory element and *LMNB1* we used a powerful, recently developed, sequence-based deep-learning approach, known as *Orca* (Fig. 5f, Fig. 6)^32^. *Orca* can predict, from sequence data, the 3D genome structure from kilobase-scale up to whole-chromosome-scale^32^. Remarkably, all ADLD disease-causing SVs (ADLD-Dup, ADLD-Del and ADLD-Inv-Dup) reduced interactions between the 19kb critical region and one of the copies of the *LMNB1* gene (Fig. 6a-e). The direction and magnitude of genomic interaction change predicted by *Orca* was quantified by the delta score (Δ), defined as the logarithmic fold change of the interaction score between the promoter region of the *LMNB1* gene and the 19 kb critical region (log Mutant/WT, see Methods section for details). Specifically, the majority of the ADLD-causing duplications (ADLD-Dup majority) resulted in a duplicated *LMNB1* promoter that had a dramatically reduced interaction with the critical 19kb region, compared to the original promoter (Δ_2_=-1.79, Fig. 6b). In the case of A8, only a portion of the 19kb region was duplicated and the duplicated *LMNB1* gene interacted only with partial regulatory element ((Δ_2_=-0.56, Fig. 6c). ADLD-del removed the critical region completely and therefore the interaction with the critical region was lost (Fig. 6d). Interestingly, for ADLD-Del, *Orca* was able to predict a novel interaction between an upstream element representing an ectopic enhancer in ADLD-del patients that had been previously identified^7^, further confirming the accuracy of the *Orca* model (Fig. 6d).

**Figure 6:**
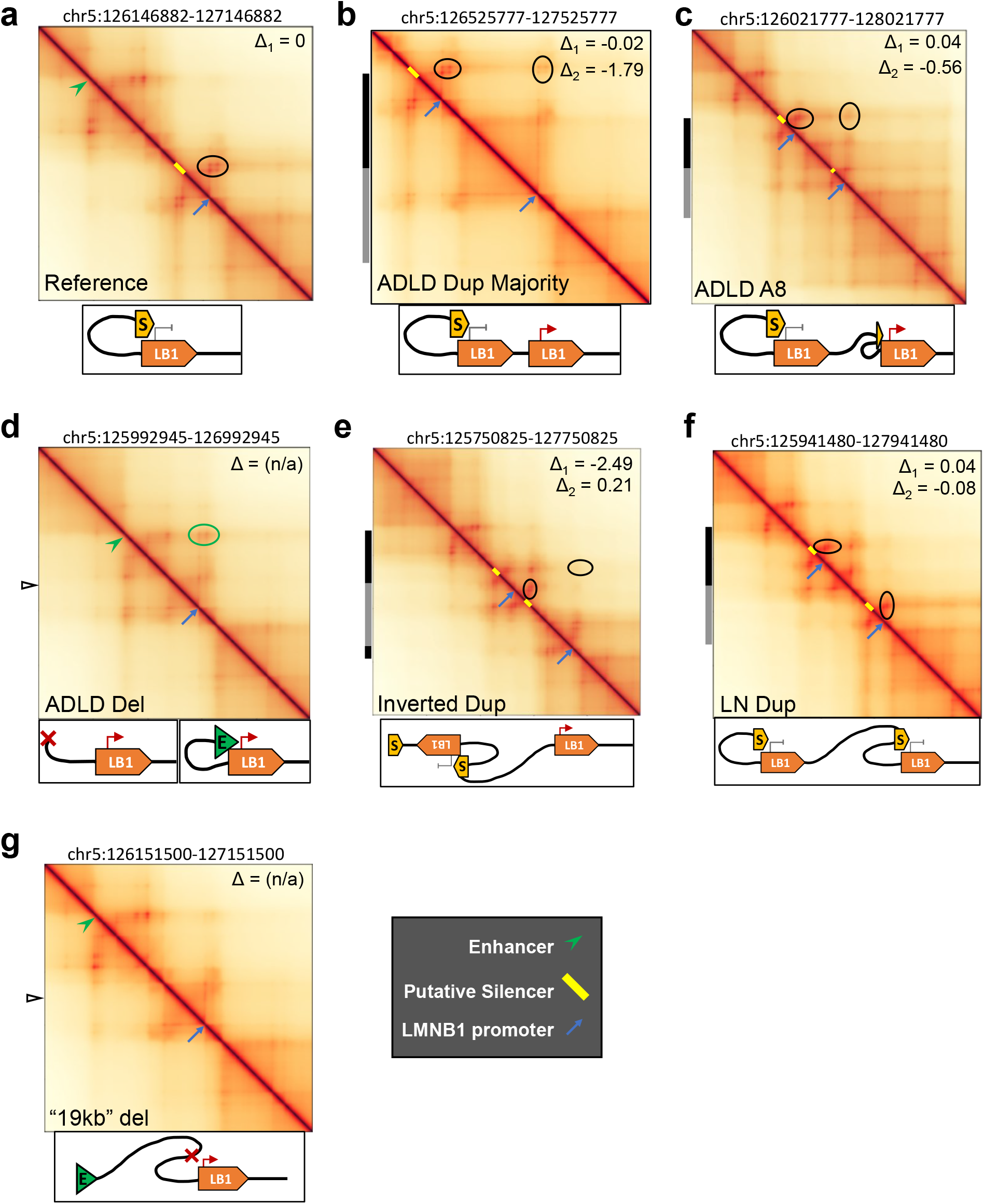
*Orca* simulation of 3D chromatin architecture in *LMNB1* structural variants. **a** *Orca* plot of reference human genome demonstrating interaction (horizontal oval) of the *LMNB1* promoter (blue arrow) with 19 kb regulatory element (yellow bar). Schematic of model describing interaction of putative silencer element with *LMNB1* is shown below the plot. Coordinates of the genomic region displayed is above the plot. **b** Plot of ADLD-Dup majority demonstrating duplication of the *LMNB1* gene but not the silencer element. Δ_1_ represents the strength of interaction of the 19 kb element and the non-duplicated *LMNB1* promoter and is unchanged while Δ_2_ represents the strength of interaction of 19 kb element with the duplicated *LMNB1* gene (vertical oval) and is reduced. This is represented in the *Orca* plots by a reduction in color intensity. The duplicated region is represented by grey and black bars on the left side of the plot. **c** The A8 ADLD-causing duplication contains the entire *LMNB1* gene and only part of the putative silencer element. This SV also results in a reduced Δ_2_ (vertical oval). In both **b** and **c**, no other genomic regions are predicted to interact with either of the copies of the *LMNB1* gene. **d** ADLD-Del results in loss of the putative silencer element and there is no interaction with the *LMNB1* promoter (Δ cannot be calculated). The genomic deletion results in the interaction (green oval) of an exogenous enhancer (green arrowhead) with *LMNB1*. Open arrowhead indicates the site of the junction subsequent to the deletion. **e** The ADLD-causing inverted duplication is inserted between the original copy of the putative silencer element and its cognate *LMNB1* gene and results in the loss of interaction between them and demonstrated by a reduced Δ_1_ (horizontal oval). A detailed schematic of the *Orca* prediction for this SV is presented in Supplementary Fig. 7. **f** LN-Dup SVs duplicate both the *LMNB1* gene and the putative silencer element, and there is no alteration in the interaction between these two elements (Δ_1_ and Δ_2_) in either of the duplicated copies (horizontal and vertical ovals). Note that no other potential regulatory genomic regions interact with either of the copies of the *LMNB1* gene. **g** Deletion of the 19 kb putative silencer element does not result in new interactions with the *LMNB1* promoter.

Although the ADLD Inv-Dup duplication includes the 19 kb region and is similar in size to one of the LN-Dups, it paradoxically results in the most severe disease phenotype. This finding can now be explained in the context of the silencer model of *LMNB1* regulation (Fig2a, Fig. 6e). The inverted, duplicated copy of the *LMNB1* gene together with its 19 kb regulatory element are inserted between the original *LMNB1* gene and its cognate 19 kb region. The *Orca*-prediction demonstrated that this inverted and duplicated 19 kb element forms a novel interaction with the original 19 kb element effectively sequestering the latter element and preventing its interaction with the non-duplicated *LMNB1* gene (Fig. 6E, Supplementary Fig. 7). As a result, the interaction of the original 19 kb element and the non-duplicated *LMNB1* gene and the Δ score are reduced to an even greater extent (Δ_1_=-2.49) than when compared the canonical ADLD duplications. One can expect that this rearrangement would lead to the overexpression of *LMNB1* to be even higher and thus lead to increased disease severity as observed in the F4-1 patient.

In contrast, the LN-Dup SVs duplicated both *LMNB1* promoter and the 19 kb critical region and the interactions for these two regions were preserved for both original and duplicated gene-silencer pairs as evidenced by a minimal alteration in the delta score (Δ_1_=0.04, Δ_2_=-0.08, Fig. 6f), consistent with the lack of an ADLD phenotype associated with this SV. For both the ADLD-Dup and LN-Dup cases, *Orca* also predicted that no new long-range regulatory elements (either enhancers or silencers) interact with the original or duplicated copies of the *LMNB1* promoter and that the loss of interactions with the 19 kb regulatory region in the ADLD-Dup is the only difference between these two SVs (Fig 6b, c, f). Orca also predicted that deletion of the 19 kb regulatory element alone did not result in the formation of any new 3D chromatin interaction with *LMNB1* (Fig. 6g). If the 19 kb region is a silencer element, as we predict, this loss of interaction in the ADLD-causing SVs should lead to an increase in *LMNB1* gene expression.

### Deletion of 19kb candidate silencer region results in oligodendrocyte specific overexpression of *Lmnb1*

To determine if the 19 kb genomic segment indeed acts as a silencer element, we deleted this region in the three mouse cell types described above and observed an increase in *Lmnb1* expression only in oli-neu cells but not in either the N2A or 3T3 cells (Fig 7a, b, Supplementary Fig. 8). The fold increase in *Lmnb1* expression was similar to the deletion of the 134 kb ADLD critical region and suggested that silencing capacity of the ADLD critical region resided within this 19 kb element. To confirm that the increase in *Lmnb1* expression was not an artifact of the CRISPR-Cas9 genomic editing process we also made a similar sized deletion outside the *Lmnb1* TAD and did not observe any differences in *Lmnb1* expression between CRISPR treated and control oli-neu, N2A and 3T3 cells (Supplementary Fig. 9). Given that the 19 kb region does not disrupt the TAD boundary containing *Lmnb1* and that the magnitude of overexpression was similar between the 134 and 19 kb deletions, it also suggests that the cause of the overexpression is unlikely due to the interaction with a novel enhancer element from outside the *Lmnb1* TAD as has been previously suggested^7^. The 19 kb deletion did not alter the expression of other genes in the nearby vicinity (*Gramd3, Aldh7a1*, and *Phax*), confirming that this regulatory element was specific for *Lmnb1* (Supplementary Fig. 8).

**Figure 7:**
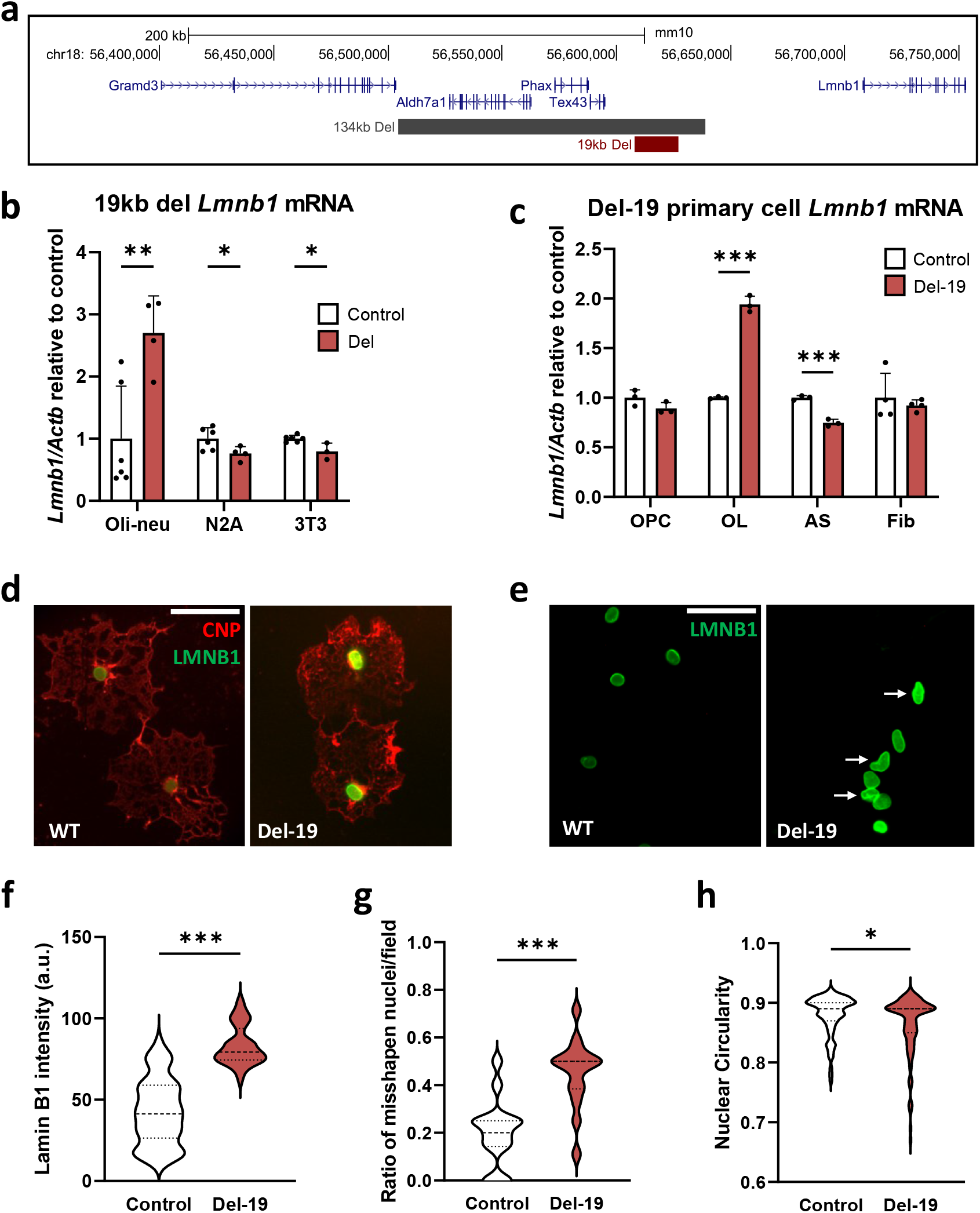
Deletion of the 19 kb regulatory element results in increased OL *Lmnb1* expression and nuclear abnormalities. **a** UCSC genome browser view of genomic segment encompassing mouse *Lmnb1* showing the 134 kb and 19 kb deleted regions. **b** Real-time PCR analysis of *Lmnb1* mRNA expression in oli-neu, N2A and 3T3 cells with the 19 kb deletion. *Lmnb1* expression is significantly higher in oli-neu cells with the deletion but lowered in N2A and 3T3 cells, relative to their respective controls. *Lmnb1* expression is normalized to beta actin (*Actb*). n ≥ 3 unique cell lines for both control and deletion clones for each cell type. *: p < 0.05, **: p< 0.01, t-test. **c** Real-time PCR analysis of *Lmnb1* mRNA expression in primary oligodendrocyte progenitor cells (OPC), oligodendrocytes (OL), astrocytes (AS), and ear fibroblasts (Fib) isolated from *Lmnb1*-Del19 (Del-19) and control mice. *Lmnb1* is significantly increased in OLs but reduced in astrocytes and unchanged in OPCs and fibroblasts. **d** Representative epifluorescence images of cultured primary differentiated OLs from Del-19 mice and WT controls stained with antibodies against LMNB1 (green) and the OL specific marker CNP (red). Scale bar = 50*μ*m. **e** Staining of OL nuclei from Del-19 mice and WT demonstrate increased presence of misshapen nuclei in the former (arrows). Scale bar = 50*μ*m. Violin plot quantifications of **f** LMNB1 intensity, **g** ratio of misshapen nuclei and **h** circularity reveal increased *LMNB1* intensity and ratio of misshapen nuclei and decreased nuclear circularity in OLs from the Del19 mice relative to control cells. n = ∼ 90 cells for each genotype. *: p < 0.05. ***: p < 0.001, calculated using two-tailed Mann-Whitney test.

To further confirm that the 19 kb region silences *Lmnb1* expression specifically in oligodendrocytes, we generated a mouse line (*Lmnb1*-Del-19) with a germline deletion of this genomic segment. The deletion was generated using the same gRNAs used to generate the deletion in mouse oli-neu cells (Supplementary Fig. 10). Primary OLs isolated from these mice exhibited a significant increase of ∼2 fold in *Lmnb1* levels compared to wild type controls, similar in magnitude to what we have observed in oli-neu cells (Fig. 7b-e). Such an increase was not observed in ear fibroblasts or primary astrocytes, representing non-OL cell types (Fig. 7c).

One of the most recognizable consequences of lamin B1 overexpression across multiple cell types is an alteration of nuclear structure, and we used this as a functional readout of lamin B1 overexpression^3,33,34^. We identified an increase in the frequency of misshapen nuclei and reduction in nuclear circularity in OL nuclei derived from *Lmnb1*-Del-19 mice when compared to wild type (Fig 7d-h), consistent with previous reports of impact of lamin B1 overexpression on nuclear architecture^33,34^. Such nuclear abnormalities were not observed in astrocytes from the *Lmnb1*-Del 19 mice (Supplementary Fig. 11).

The results from the mouse model further strengthen our OL-specific silencer mediated *Lmnb1* overexpression hypothesis and confirm the results we obtained in the oli-neu cell lines.

### CTCF binding sites within the 19 kb region mediate *Lmnb1* silencing

To elucidate the molecular mechanisms underlying the silencer element we carried out a bioinformatic analysis of transcription factor bindings sites in the two regions within the 19 kb element that interact with the lamin B1 promoter in both mouse and human genomes. In humans, we observed that both regions were enriched for binding by the transcription factor CCCTC binding factor (CTCF) that we named CTCF1 (chr5: 126656590-126656600) and CTCF2 (chr5:126676270-126676280) (Supplementary Fig. 12a). Based on the Hi-C data (Fig. 5a) these CTCF binding sites interact with CTCF binding sites CTCF3 (chr5:126780430-126780440) and CTCF4 (hg38 - chr5:126676270-126676280) in the first and third introns of the *LMNB1* gene. The CTCF1 (mm10-56608876-56608895), 2 (mm10-56622463-56622482) and 3 (mm10-56711264 - 56711282) sites were also conserved in the mouse genome while CTCF 4 was not (Supplementary Fig. 12b). Within TADs, one of the main functions of CTCF is to target regulatory elements to their cognate promoters by forming chromatin loops and loop forming CTCF sites need to be in opposite orientations^35–37^. We observed that CTCF1 and CTCF2 were in the forward orientation while CTCF3 and CTCF4 were in the reverse orientation confirming their ability to from a chromatin loop as indicated by the Hi-C and *Orca* maps (Supplementary Fig. 12c).

We confirmed that the CTCF 1, 2 and 3 predicted sites were indeed bound by the CTCF protein by carrying out genome wide analysis of CTCF binding in primary human fibroblasts and mouse embryonic stem (ES) cells and primary oligodendrocytes using the Cleavage Under Targets and Release Using Nuclease (CUT&RUN) technique^38,39^ (Supplementary Fig. 13, Fig. 8a-c). The conservation of CTCF binding across cells types and species would indicate that they have a functionally important role in lamin B1 regulation.

**Figure 8:**
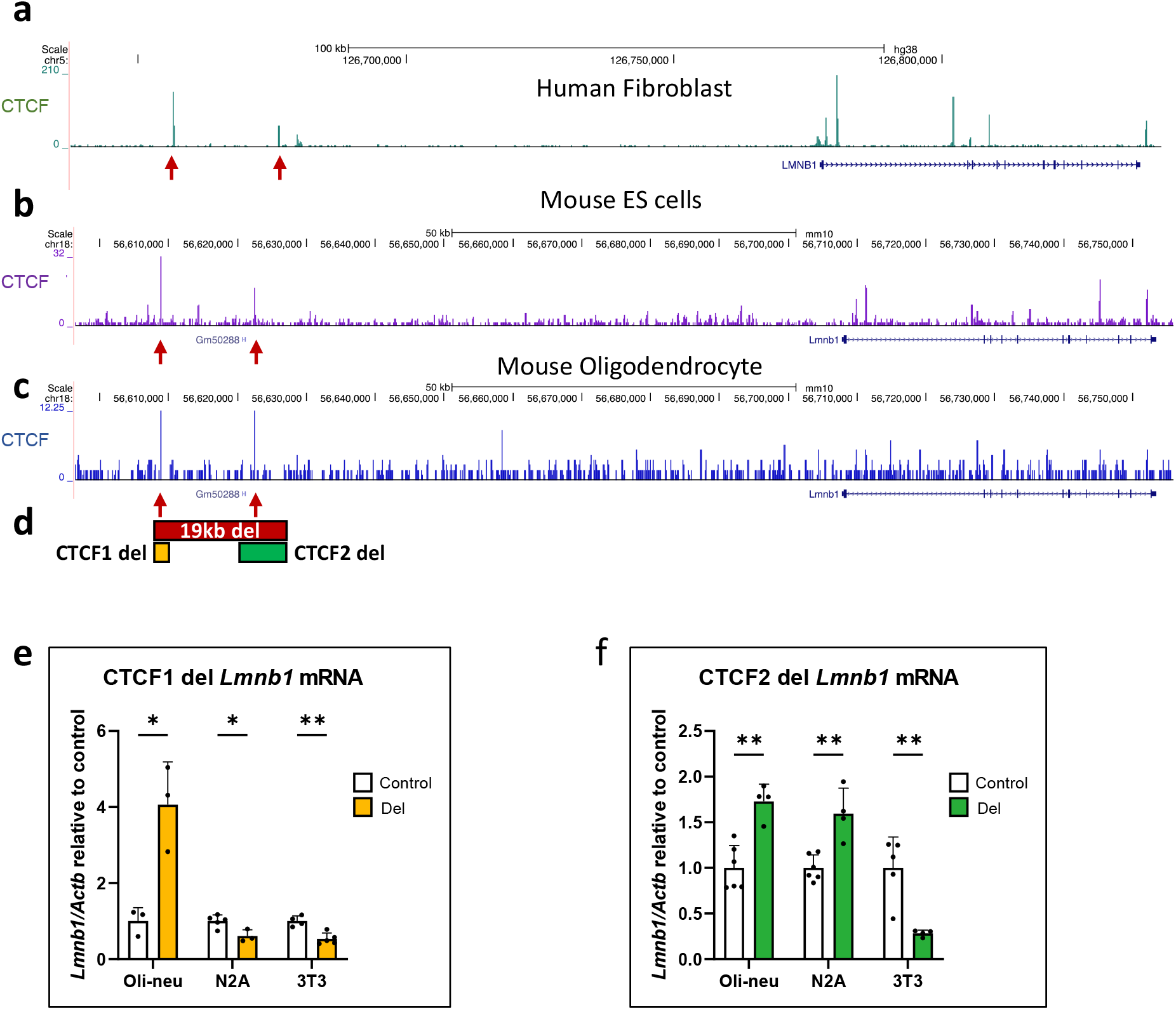
Identification of CTCF binding in 19 kb silencer element and consequences of CTCF binding site deletion. UCSC genome browser tracks of CUT&RUN analysis in **a** human fibroblast, **b** mouse ES cells and **c** mouse primary oligodendrocytes confirms bioinformatically predicted CTCF binding sites within 19 kb silencer element. Arrows point to the conserved CTCF 1 and CTCF 2 sites. **d** Schematic showing deletions of CTCF 1 and 2 sites. Real-time PCR analysis of *Lmnb1* mRNA expression in oli-neu, N2A and 3T3 cells with CRISPR-mediated deletions of **e** CTCF1 and **f** CTCF2. *Lmnb1* expression is significantly higher in oli-neu cells with CTCF1 deleted but lowered in N2A and 3T3 cells, relative to their respective controls. *Lmnb1* expression is increased in oli-neu and N2A cells with CTCF2 deleted but lowered in 3T3 cells. *Lmnb1* is normalized to b Actin (*Actb*). Graphs are mean ± SEM and n ≥ 3 unique cell lines for both control and deletion clones for each cell type. *: p < 0.05, **: p < 0.01, calculated using Student’s t test.

To experimentally test whether the individual disruption of either site impacted *Lmnb1* expression, we generated CRISPR deletions that removed each CTCF site independently in oli-neu, N2A and 3T3 cell lines (Supplementary Figs. 14, 15). Deletions of the CTCF1 site resulted in an increase in *Lmnb1* expression in oli-neu cells but not in the other two cell types while deletion of the CTCF2 site resulted in increased *Lmnb1* expression in oli-neu and N2A cells but not in 3T3 cells (Fig. 8d, e). The magnitude of the increase was similar to the deletion of the 19 kb critical region indicating that these sites were responsible for the silencer function of the 19 kb region.

To further explore the role of CTCF in mediating *Lmnb1* silencing in OLs, we carried out RNAi mediated knockdown of CTCF in all three cell types and observed an increased expression of *Lmnb1* only in oli-neu cells but not in N2A or 3T3 cells (Fig. 9b, Supplementary Fig. 16a). Consistent with our model, a similar experiment using the oli-neu cells with the 19 kb deletion did not exhibit any increase in *Lmnb1* expression when treated with CTCF RNAi (Fig. 9b), confirming that CTCF binding specifically in the 19 kb critical region was responsible for transcriptional downregulation of *Lmnb1*.

**Figure 9:**
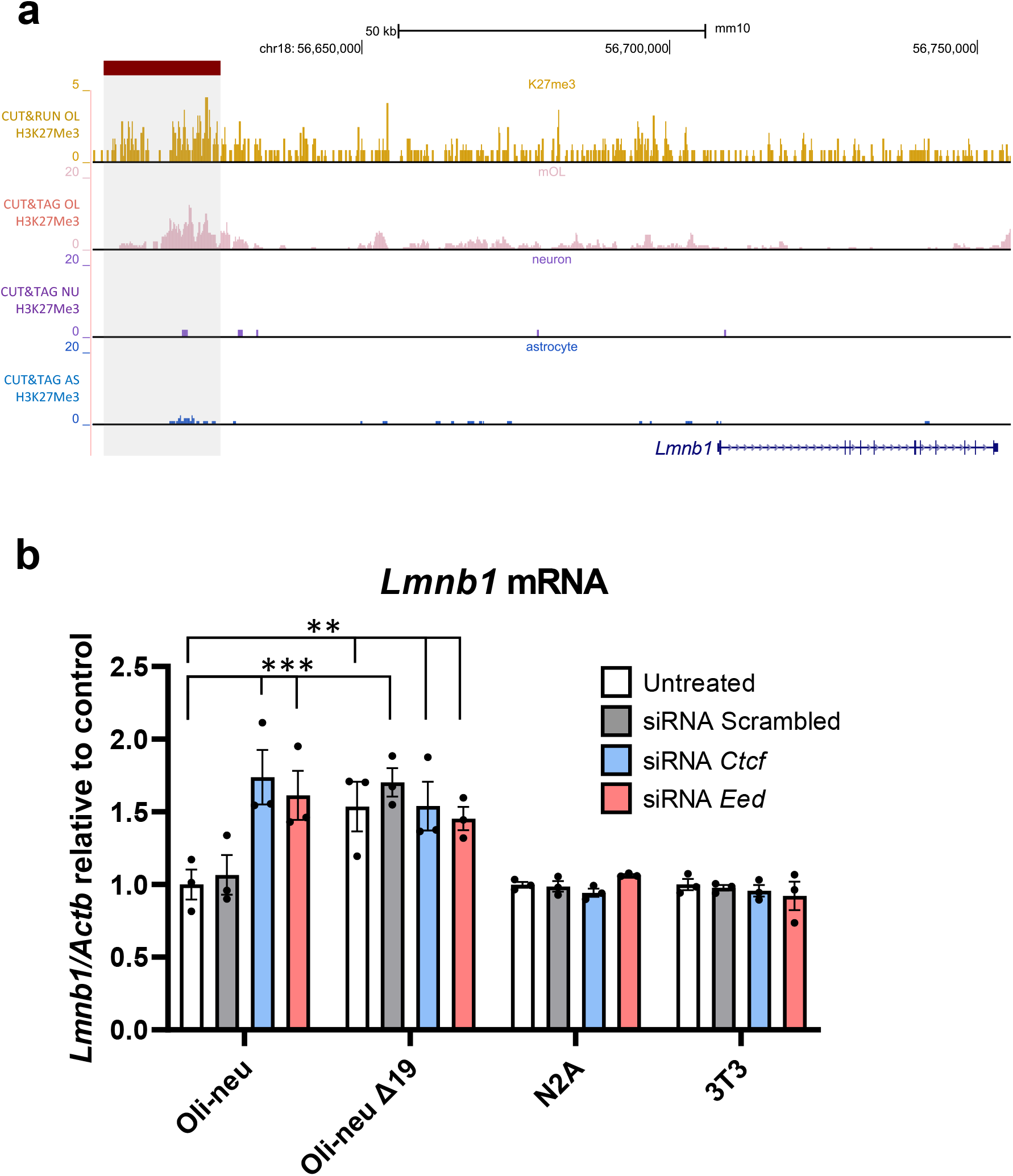
Epigenetic analysis and CTCF and PRC2 complex involvement in OL specific *Lmnb1* silencer element. **a** Genome tracks of H3K27Me3 CUT&RUN analysis of mouse OLs and CUT&TAG analysis of mouse OLs, astrocytes (AS) and neurons (NU) reveal enrichment of H3K27Me3 in the 19kb silencer (grey shaded region) element specifically in OLs. CUT&TAG analysis is from previously published data^25^. **b** *Ctcf* and *Eed* siRNA treatment of oli-neu, N2A and 3T3 cell lines reveals increased *Lmnb1* expression only in oli-neu cells, relative to respective untreated cell specific controls. Scrambled siRNA treatment has no effect on *Lmnb1* expression. No increase in *Lmnb1* expression is observed in oli-neu cells with the 19 kb deletion (oli-neu Δ19) relative to untreated Δ19 cells. Graphs are mean ± SEM for 3 biological replicates per cell type and treatment. ***: p < 0.001, using two-way ANOVA.

### Epigenetic analysis is of the 19kb region identifies a role for the PRC2 complex in lamin B1 silencing

One mechanism by which CTCF mediates transcriptional silencing is by its association with the polycomb repressive complex 2 (PRC2) group of proteins^40,41^. The PRC2 is a large multimeric protein complex and has been shown to function as a repressor protein complex in the establishment of long-range chromatin interactions^42,43^. The PRC2 complex comprises of the core subunits, SUZ12, EED, and the methyltransferase EZH2 or its closely related homolog EZH1, and are all required for the catalytic activity of the complex^44^.

Chromatin regions that are silenced by the PRC2 complex are enriched in H3K27 tri-methylation (H3K27Me3)^43,45^, and we analyzed previously published data on H3K27Me3 binding within the *Lmnb1* TAD from various mouse CNS cell types^25^, and also carried out H3K27Me3 CUT&RUN analysis on primary OLs (Supplementary Fig. 17). Strikingly, we observed a significant enrichment of H3K27Me3 binding within the 19 kb silencer region only in OLs but not in other CNS cell types such as astrocytes and neurons (Fig. 9a).

Similar to CTCF, we carried out RNAi mediated knockdown of one of the PRC2 components, *Eed*, in all the three cell types and observed increased expression of *Lmnb1* only in oli-neu cells but not in N2A or 3T3 cells (Fig. 9b. Supplementary Fig. 16b). As predicted, *Eed* knockdown in the 19kb del oli-neu cells did not exhibit any further increase in *Lmnb1* levels (Fig. 9b). Supporting our findings, an examination of previously published RNA-Seq data revealed that *Lmnb1* expression was significantly increased by ∼1.5 fold compared to controls in mouse OLs where the *Eed* was conditionally knocked out^46^.

## Discussion

Our results provide insights into the tissue specificity of ADLD and provide a unifying mechanism to explain how duplications and upstream deletions can cause disease. They indicate that ADLD is caused by a transcriptional targeting of lamin B1 overexpression specifically to OLs, mediated by the loss of an interaction with an OL specific silencer element, rather than just the presence of an extra copy of the *LMNB1* gene.

These results also establish a framework for the interpretation of the pathogenic potential of *LMNB1* SVs, and other pathogenic SVs in general, that have significant implications for genetic testing and provide insights into how the disruption of long-range regulatory elements can lead to disease. For the first time, our findings demonstrate that not all *LMNB1* duplications will result in ADLD, and we indicate that the size, location and insertion site of the *LMNB1* duplications are all essential for predicting disease onset. Loss of interactions with the silencer element appear to be the critical determinant of pathogenicity of the *LMNB1* SVs (Supplementary Fig. 18). In the case of ADLD-Del, it is yet unclear if interaction with the ectopic enhancer is also required for disease causation or whether deletion of the silencer element alone is sufficient. While our results favor the latter hypothesis, loss of the silencer element would still be required to facilitate the interaction of the ectopic enhancer with the *LMNB1* promoter and would thus be a critical first step in ADLD pathogenesis (Supplementary fig. 18).

Once the exact location, size and orientation of the SVs are known, interaction with the silencer element can be predicted using bioinformatics tools such as *Orca*. While this is straightforward for deletions that remove the silencer element, this can be more for complicated for duplications or inversions and a case by case review is required to predict pathogenicity. For simple tandem duplications that encompass the silencer element, we would predict that ADLD would not develop. However, if the duplications are not tandem, it is critical to identify their exact insertion site as they may be inserted in a manner that disrupts the lamin B1-silencer interaction, as has occurred in the F4-1. While patients with the large duplications do not exhibit symptoms of ADLD, some do exhibit more subtle clinical phenotypes including those of autonomic dysfunction, and the exact role of *LMNB1* duplications in these cases needs to be explored further.

When predicting the pathogenic impact of SVs, conventional wisdom has dictated that the larger the variant and the more genes involved, the greater likelihood that the variant will be deleterious^47^. Paradoxically, our results demonstrate that the larger duplications have a more benign impact and provide a rationale for explaining this phenomenon that may also be applicable to other disease-causing SVs. There have been very few reports implicating silencer elements in disease pathogenesis, especially those that silence target gene expression in specific cell types and none for Mendelian diseases^48^. To the best of our knowledge, we have also not encountered previous reports of duplications leading loss of interactions with a silencer element as a disease mechanism. These represent novel consequences of genomic alterations that should be borne in mind with predicting the pathogenic potential of other SVs.

Our results indicate that OLs are the primary cell type that are targets of the lamin B1 overexpression and therefore the primary cell type that drives the ADLD disease process. This finding is consistent with our previous reports that transgenic mice where *LMNB1* overexpression is targeted to OLs exhibit ADLD-like phenotypes, while targeting *LMNB1* overexpression to astrocytes or neurons does not result in demyelination^13,14^. Histopathological analysis of ADLD brain post-mortem tissue revealed no alteration in OL number but has identified abnormally shaped astrocytes as a hallmark of the disease^15,16,49^. Given that we did not identify overexpression of lamin B1 in astrocytes from the *Lmnb1*-Del-19 mice, this would indicate that astrocyte dysmorphology is likely a secondary, cell non-autonomous consequence of OL dysfunction. The finding that OLs are the primary driver for the ADLD disease process also have important implications for designing therapeutic approaches, especially for those that utilize lamin B1 reduction as a strategy. These approaches can now be targeted specifically to OLs and can reduce the potential side effects of a more indiscriminate reduction of lamin B1 levels in other cell types such as astrocytes or neurons.

Our findings are consistent with a recent report that has characterized a class of silencer elements known as H3K27me3-rich regions or methylation rich regions (MRRs)^45^. As we have demonstrated for lamin B1, these MRRs are located at sites of long-range chromatin interactions and are thought to function through chromatin looping. They also answer an important question of whether the silencing effects are directly mediated by PRC2, or whether PRC2 is an effector of CTCF–cohesin-mediated chromatin looping^48^. Our results suggest the latter as the chromatin loops linking the silencer element are conserved across cell types while H3K7Me3 modifications are only observed in OLs. This suggests that the PRC2 complex is specifically recruited to the silencer element only in OLs indicating that PRC2 is not required for the formation of the CTCF-mediated chromatin loops.

A yet unanswered question that arises from our findings is the mechanism underlying the OL specificity of the silencer we have identified. How this occurs is unclear and would likely involve the interaction of an OL specific transcription factor with PRC2. While enhancers in OLs have been previously reported^20,21^, this is the first report of an OL specific silencer element. Our results can help identify additional OL specific silencer elements and determine whether CTCF-PRC2 interactions are a common mechanism for the downregulation of other genes in maturing OLs. They also provide insights into how long-range non-coding regulatory element can modulate gene expression and identify a hitherto unknown role for silencer elements in tissue specificity and disease causation.

## Materials and Methods

### Clinical and MRI examination and collection of tissue samples

Clinical and magnetic resonance imaging (MRI) examination of all subjects were carried out at the respective clinical centers after appropriate consent and IRB approval. Fibroblast samples were obtained from skin punches and cultured as described below. Brain tissue was collected from ADLD patients at autopsy and samples were flash frozen. In all cases, affected frontal white matter and unaffected frontal grey matter was used for RNA isolation. Control brain samples were obtained from same brain region from age and sex matched individuals.

### Genomic DNA isolation and Array-based comparative genomic hybridization (Array CGH)

Genomic DNA from whole blood, saliva, primary fibroblasts and brain tissue was isolated using the Gentra Puregene kit (Qiagen) according to the manufacturer’s instructions. Array CGH was performed at the University of Pittsburgh on genomic DNA from clinical samples, hybridized on a custom 8×15K HD-CGH microarray previously described^5^, scanned in an G2565CA Agilent microarray scanner, and analyzed using Agilent CGH Analytics software (Agilent Technologies). Human genome assembly GRCh38/hg38 was used for all genome coordinates. PCR primers were designed to specifically amplify the tandem duplication junctions for each family using Longamp Taq Polymerase (New England Biolabs), then Sanger sequenced (Eurofins Genomics) after treatment with ExoSAP-IT reagent (Applied Biosystems). The duplication junction primers sequences are listed in Supplementary table 2

### CRISPR guide RNA design, generation and validation of CRISPR clones

“Left” and “right” CRISPR guide RNA (gRNA) sequences for either ends of the genomic deletions were selected using the CRISPOR online tool^50^, selecting for high specificity and efficiency, adjacent to *S. pyogenes* Cas9 Protospacer Adjacent Motif (PAM) sites (NGG), encompassing the regions of interest in the mouse genome (mm10) on chromosome 18. The genomic targets of the guides were Sanger sequenced to ensure there were no known variants in the binding sequences or PAM. Both guides were cloned into pDG458 (Addgene) via golden gate cloning, as previously described^51^, then purified using EndoFree Plasmid Maxi Kit (Qiagen). The gRNA spacer sequences for each of the deletions are listed in Supplementary table 2.

The purified dual-guide CRISPR plasmids were transfected into oli-neu, N2A, and 3T3 cells using Lipofectamine 3000 (ThermoFisher), grown for 24 hours post-transfection. GFP-positive cells were than sorted using Fluorescence Activated Cell Sorting (FACS) as single cells into 96-well plates to establish clonal populations and subsequently genotyped for deletions using DirectPCR Cell Lysis buffer (Viagen), and cells positive for deletions were expanded (Fig. 4c).

For clone validation, genomic DNA was isolated from cells using Gentra Puregene Kit (Qiagen) and CRISPR-induced deletions were confirmed via PCR using primers adjacent to each end of the deletion. The primers to sequence deletion junctions are as listed in Supplementary table 2. Amplicons were Sanger sequenced (Azenta) to verify the coordinates and uniqueness of deletions. Clones with deletions were used for expression analysis via quantitative RT-PCR. Cell lines were also tested to ensure they did not contain genomic inversion via PCR using a three-primer strategy as described in Supplementary Figure 6c.

### Cell culture

For fibroblast isolation, patient and control skin biopsies were minced to ∼1.0 mm^3^ pieces then incubated with 0.05% Trypsin-EDTA (Sigma) for 3 hours at 37°C with gentle mixing. Pieces were then plated on a 10-cm tissue culture dish (ThermoFisher) with a minimal amount of DMEM complete (high glucose DMEM (Corning) supplemented with 10% FBS (Fisher), 2mM L-glutamine (Millipore), and 1% penicillin-streptomycin (Hyclone)) until they adhered to the culture dish. Primary fibroblasts were trypsinized and re-plated into new dishes as they exited the explant. N2A and NIH-3T3 were cultured grown in DMEM complete medium. Oli-neu cells were cultured as described previously^52^. Astrocytes and OPCs were isolated by immunopanning from 6-7 day old mouse pups and OPCs were differentiated into OLs for 4 days according to previously-established protocols^53^. All cells were cultured at 37°C and 5% CO_2_ in a humidified chamber.

### RNAi treatment

For transfection of RNAi constructs, cells were seeded at a density of ∼1 × 10^5^ cells per well of a six-well plate for 16–18 hours prior to transfection. 50-75 pmol of pre-designed MISSION® esiRNA (Sigma) using Lipofectamine™ RNAiMAX Transfection Reagent (Thermo Fisher) were transfected in each well. After 24 hours the medium was replaced with fresh proliferation medium and cultured another 24 hours. In the case of oli-neu cells, they were then switched to differentiation medium for 4 days.

### Generation of *Lmnb1*-Del-19 mice

These mice were generated using CRISPR/Cas9 rechnology as previously described^54^. Briefly guide RNAs used to generate the 19 kb deletion and cloned in the pDG458 plasmid were *in-vitro* transcribed using the TranscriptAID T7 High Yield Transcription Kit (ThermoFisher) and purified using GeneJET RNA purification kit (ThermoFisher). Guide RNAs (200 ng/μl) together with the Cas9 protein (100 ng/μl, Alt.R S.p. Cas9) Nuclease 3NLS, IDT) were electroporated into C57B6 mouse zygotes and transferred the following day as two-cell stage embryos into the oviducts of pseudo-pregnant females. Founder lines were screened using primers designed to amplify across the 19 kb deletion listed in Supplementary table 2.

### RNA isolation, cDNA synthesis, and real-time PCR

RNA was isolated from cells, sciatic nerves and brain tissue using TRIzol reagent (Invitrogen) following the manufacturer’s instructions. cDNA was synthesized from 1 μg of RNA using qScript cDNA Synthesis Kit (Quanta Bio). Real-time PCR was performed using PerfeCTa SYBR Green SuperMix with ROX (Quanta Bio) on an ABI QuantStudio 12K Flex (Applied Biosystems). Gene expression was analyzed using the ΔΔC_T_^55^ method using *Actb* mRNA as an endogenous control. Primer efficiencies were validated using 6-fold serial dilutions of cDNA. Primers used for real time PCR analyses are listed in Supplementary table 2.

### Real-time copy number analysis

Purified genomic DNA samples from control and CRISPR-edited cells were diluted to 5 ng/µL and real-time PCR was performed using PerfeCTa SYBR Green SuperMix with ROX (Quanta Bio) on an ABI QuantStudio 12K Flex (Applied Biosystems). Copy numbers were calculated using the ΔΔC_T_ method^55^ normalized against *Actb* genomic DNA and plotted as relative to undeleted control lines across three regions: within the deletion region and on each side (centromeric or telomeric) of the deleted region. Cell lines without a significant reduction of deleted region DNA abundance compared to undeleted control were omitted from analysis. Copy number analysis real-time primer sequences are listed in Supplementary table 2.

### Immunocytochemisty (IHC)

IHC was performed as described previously^11^. Primary antibodies used and dilutions are listed in Supplementary table 3. AlexaFluor 488 or Cy3-conjugated secondary antibodies were used (Jackson ImmunoResearch) After antibody staining, cells were mounted onto glass microscope slides with Vectashield antifade mounting medium containing DAPI (Vector Laboratories) and imaged using a Leica CTR5000 fluorescence microscope using identical exposure settings.

### Protein isolation and Western blotting

Sciatic nerves were harvested from WT and *PLP*-*LMNB1* (TG) mice and stored at −80°C until protein isolation. To isolate protein, nerves were incubated in T-PER supplemented with 1x protease inhibitor cocktail (Thermo Scientific) and homogenized. Protein isolation wand Western blotting were carried out as described previously with 50µg protein loaded per well on a 10% gel^11^. The blot was imaged and quantified using the LI-COR Odyssey CLx infrared scanner and Image Studio software (LI-COR Biosciences). Antibodies used are listed in Supplementary Table 3.

### Myelin Visualization and Electron Microscopy analysis

To assess myelin in the periphery, sciatic nerves from 9-month-old WT and *PLP-LMNB1* mice were stained with Luxol fast blue (LFB) as previously described^56^. Sections were examined and images captured using a Leica CTR5000. Staining was quantified from five representative areas per nerve using ImageJ and an average mean intensity was measured for each nerve.

For transmission electron microscopy (TEM) analysis, mice were perfused with cold PBS and 4% PFA, sciatic nerves from WT and *PLP-LMNB1* mice were post-fixed in cold 2.5% glutaraldehyde in 0.01 M PBS (Fisher, Pittsburgh, PA), pH 7.3 and processed as previously described^13^. Sections were examined on a JEOL 1011 transmission electron microscope (JEOL Peabody, MA) or JEOL 1400 transmission electron microscope with a side mount AMT 2k digital camera (Advanced Microscopy Techniques, Danvers, MA). Images of transverse sciatic nerve sections were captured at 5,000x magnification. Counts of myelinated axons were calculated from calibrated TEM images using ImageJ.

### Sciatic nerve conduction velocity

Compound action potential (CAP) recordings were performed as previously described^57^. Briefly, Sciatic nerves were harvested from 9-month-old WT and TG mice and immediately placed in oxygenated Krebs solution. CAPs were measured at room temperature with oxygenated Krebs solution perfused by gravity into the recording chamber containing the nerves. Current was delivered through a stimulus electrode suctioned to one end of the nerve while recordings were taken from a recording electrode suctioned to the other end of the nerve. Conduction velocity was calculated by diving the length of the nerve (distance from one electrode to the other) by the latency between the initiation of stimulus artifact and peak of CAP.

### Predicting ADLD structural variant effects on 3D genome with Orca sequence models and identification of CTCF sites

We generated 3D genome interaction map predictions with Orca30 for all SVs including ADLD-dup, LN-Dup, ADLD-del, ADLD-inv-dup. As the model output values represents log fold over distanced-based expectation, we converted the prediction to log balanced count matrix scale by multiplying the model output by distance-based expectation matrix. The 3D genome predictions for both the wildtype and mutated sequences were visualized as heatmaps.

After we made the prediction for the WT and mutant sequences, we compared the interaction strength between *LMNB1* promoter and 19kb critical region in the mutant and WT samples. In the case of duplication events, we compared the interaction between duplicated critical region and the LMNB1 gene promoter with interaction strength between original copies of these two regions in the WT samples. The effects of the SV on the 3D genome interaction changes were quantified based on Orca predicted log-fold change (Δ) in interaction scores (for H1-ESC cell, Orca-32Mb predictions at 2Mb scale). When the genomic region(s) of interest were duplicated by the SV, interaction log-fold change scores for each duplicate were computed separately. CTCF sites in the 19 kb critical region and in the lamin B1 gene vicinity were identified using the Find Individual Motif Occurrences (FIMO) online tool^58^.

### CUT&RUN analysis

CUT&RUN was performed as previously described under native conditions, using recombinant Protein A-MNase (pA-MNase)^59,60^. After separating released fragments through centrifugation, fragments isolated were used as input for a library build consisting of end repair and adenylation, NEBNext stem-loop adapter ligation, and subsequent purification with AMPure XP beads (Beckman Coulter). Barcoded fragments were then amplified by 17 cycles of high-fidelity PCR and purified using AMPure XP beads. Libraries were pooled and sequenced on an Illumina NextSeq2000 to a depth of ∼10 million mapped reads. Antibodies used for CUT&RUN are listed in Supplementary table 3 CUT&RUN data was analyzed as previously described^59,60^. Paired-end FASTQ files were mapped to the mm10 genome with bowtie2 (options -q -N 1 -X 1000 --very-sensitive-local)^61^. Mapped reads were filtered for PCR duplicates using Picard tools ([http://broadinstitute.github.io/picard/) and filtered for MAPQ ≥ 10 using SAMtools^62^. Fragment distribution plots were generated using Picard. Fragment classes corresponding to TF footprints (<120 bp) were generated using a custom awk script and SAMTools. UCSC files were generated from size classed sam files using HOMER^63^. Reads were converted to bigWig files using deepTools with RPGC read normalization^64^. Heatmaps were generated using deepTools computeMatrix (options -a 2000 -b 2000 -bs 20 --missingDataAsZero) and plotHeatmap.

### List of online resources utilized

UCSC genome browser (hg38 version) - https://www.genome.ucsc.edu/

CRISPOR - http://crispor.tefor.net/

HiGlass Hi-C visualization tool - https://higlass.io/

FIMO motif finding tool - https://meme-suite.org/meme/tools/fimo

Picard tools - http://broadinstitute.github.io/picard/

BioRender - www.biorender.com

### Data availability

Additional data is provided in the Supplementary information. Detailed clinical descriptions of subjects are available upon request.

## Acknowledgements

QSP would like to acknowledge the Padiath lab members and Dr. Svetlana Yatsenko for helpful discussions and ADLD patient families for their continuing support. The work was funded by NIH grants R01NS126193, R21NS131906, R33NS104384, R33NS106087, R01NS095884 and ADLD Center grant ADLD-23-001-02 to QSP, NIH grant R35GM133732 to SJH, Clinical Research Scholar Junior 1 Award from the Fonds de Recherche du Quebec-Santé the New Investigator Salary Award from the Canadian Institutes of Health Research and the Clinical Research Scholar Senior award from the FRQS to GB. FGL holds a tier 1 Canada research Chair in tissue-engineering and in vitro brain diseases modelling. KRS and PFC are supported by the MRC International Centre for Genomic Medicine in Neuromuscular Disease (MR/S005021/1). JvdA is supported by a Wellcome Clinical Research Career Development Fellowship (219615/Z/19/Z). PFC is a Wellcome Principal Research Fellow (212219/Z/18/Z), who receives support from a Wellcome Collaborative Award (224486/Z/21/Z) and an MRC research grant (MR/S035699/1). JvdA and PFC acknowledge core funding from the MRC to the MRC Mitochondrial Biology Unit (MC_UU_00028/8 and MC_UU_00028/7). This research was supported by the NIHR Cambridge Biomedical Research Centre (BRC-1215-20014).

## Author Contributions

BN, GRB and QSP conceived and designed the overall study with contributions from SH, MG and JZ. BN, GRB, SL, TO, AJ, NH, SR and ELA carried out the experiments. KD and JZ carried our bioinformatic and *Orca* analyses. DK and YP carried out bioinformatic analysis and provided conceptual input. JR, EK, NG, KS, JvdA, PFC, SBS, FP, CT, FV, JS, LP carried out patient clinical analysis and provided patient biological material. GB and RR carried out interpretation of clinical and MRI results. JK, MT, RH and FG-L provided ADLD brain samples. QSP and BN wrote the paper with contributions from GB, KD, JZ and RR.

## Additional information

The views expressed are those of the author(s) and not necessarily those of the NIHR or the Department of Health and Social Care. For the purpose of open access, the authors have applied a Creative Commons Attribution (CC BY) license to any Author Accepted Manuscript version arising from this submission.

---

**Supplementary Figure 1:**
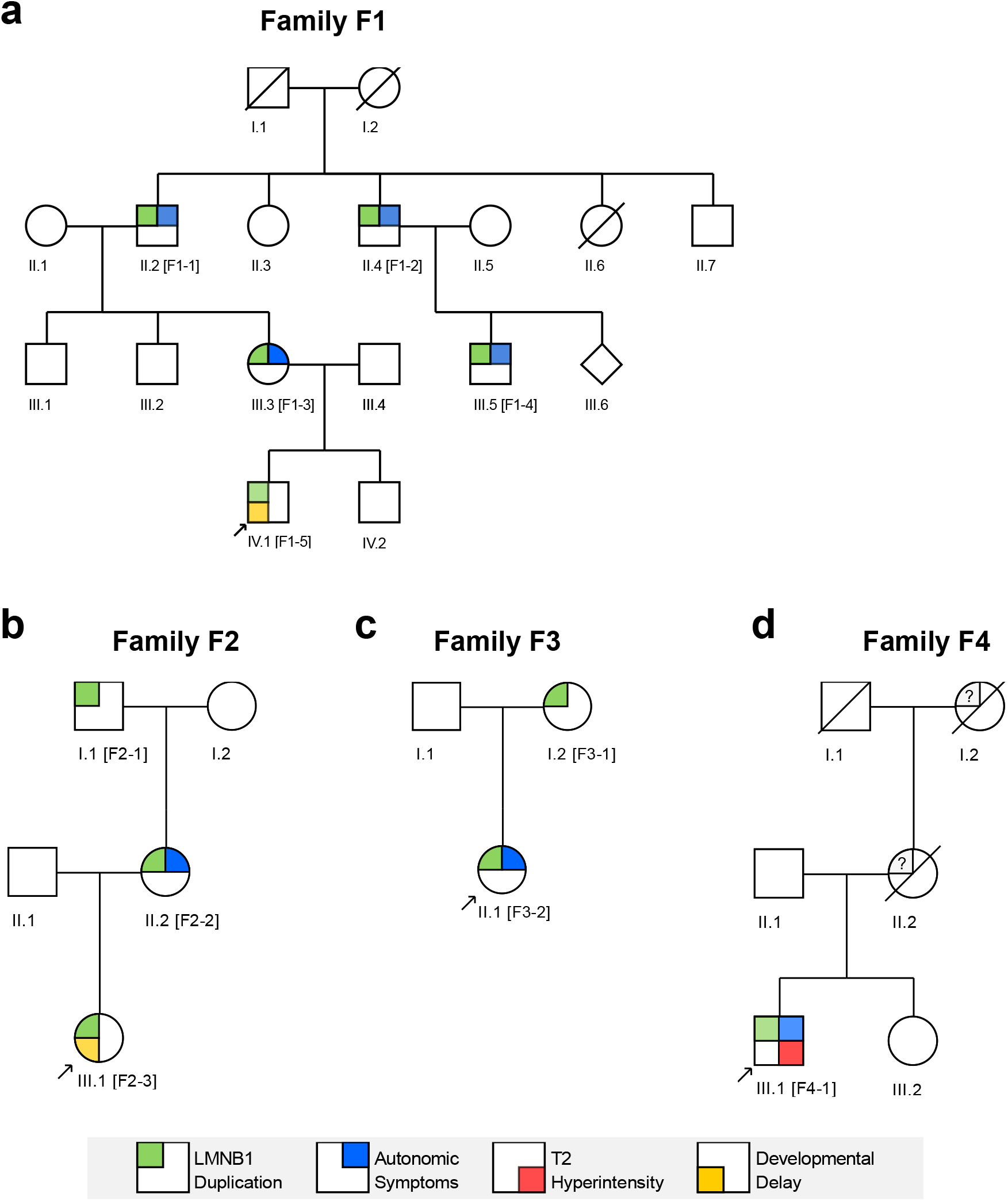
Pedigree charts of families with *LMNB1* duplications. **a-c** Families F1, F2, and F3 have duplications of *LMNB1* without the ADLD disease phenotype. **d** Subject F4-1 of family F4 exhibited a more severe ADLD phenotype. Arrows represent probands for each family. For all individuals with filled in symbols, aCGH and detailed clinical analysis were carried out with the exception of F3-1, on whom an MRI and a detailed clinical analysis was not performed. In family F4, only anecdotal information was available for members I.2 and II.2 and these are represented by question marks. Clinical details of all analyzed individuals are provided in Supplementary clinical data.

**Supplementary Figure 2:**
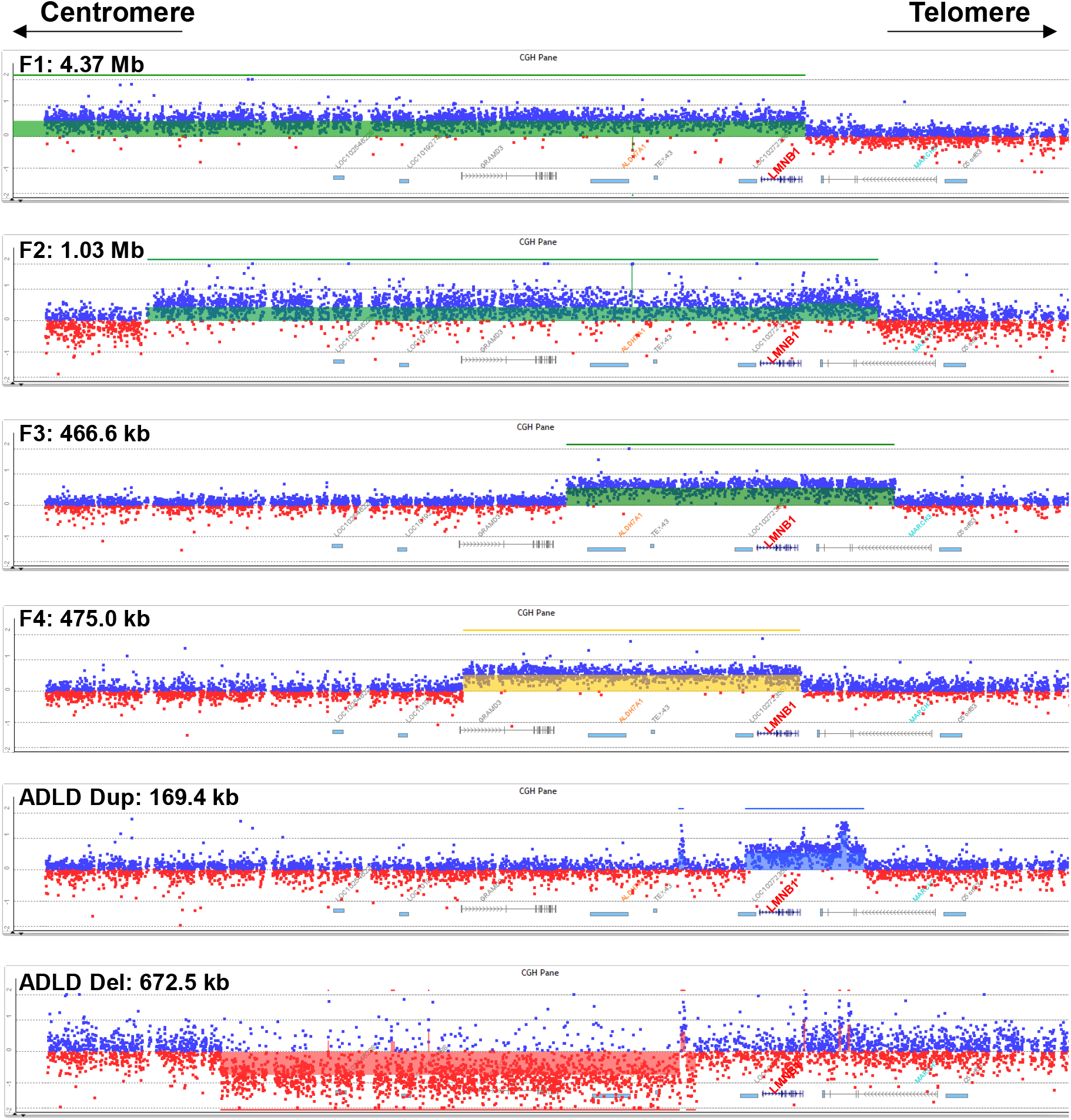
aCGH of *LMNB1* duplications. aCGH of the *LMNB1* locus using a custom designed high resolution array^5^ reveals the duplication and deletion extents in the different families. For families, F1-F3, duplications are marked by the green shaded area and by the yellow shaded area in F4. Note that for F1 the complete extent of the duplication is not shown as the custom array only extends for 2 Mb on either side of the *LMNB1* gene, and the F1 duplication extends beyond this on the centromeric side. An example of a canonical disease causing ADLD duplication and deletion are also shown for comparison (blue and red shared regions, respectively). The *LMNB1* gene is annotated in red and the Y axis represents log2 ratios.

**Supplementary Figure 3:**
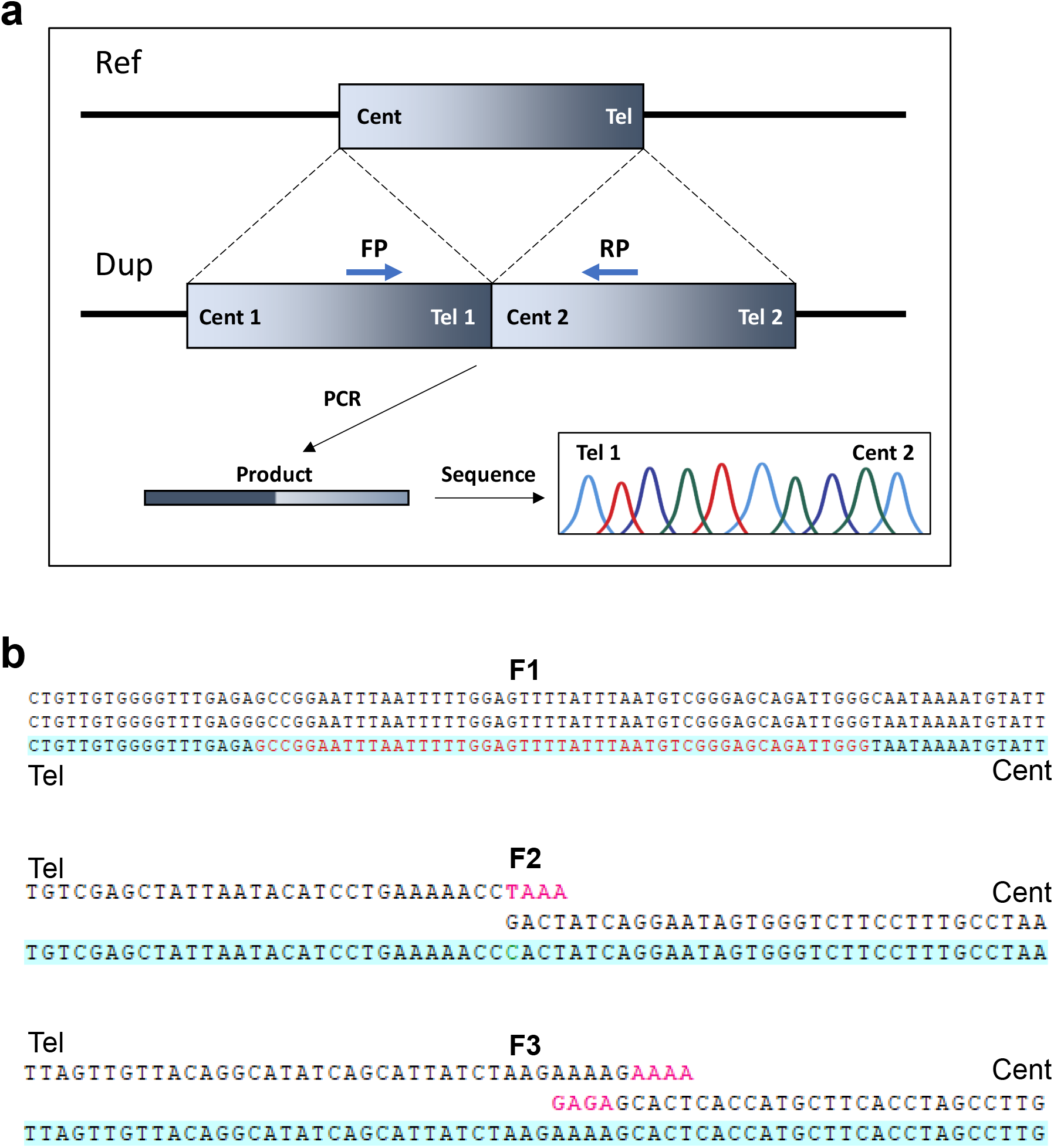
Identification of duplication junction boundaries in LN-Dup patients. **a** Schematic showing orientation of duplications and rationale for using PCR primers that flank the duplication junction to identify precise breakpoints. **b** Junction sequences for each of the LN-Dup duplications. Further details are also provided in Supplementary table 1. Note that the duplication junction for F4 has been previously described^5^ and is complex.

**Supplementary Figure 4:**
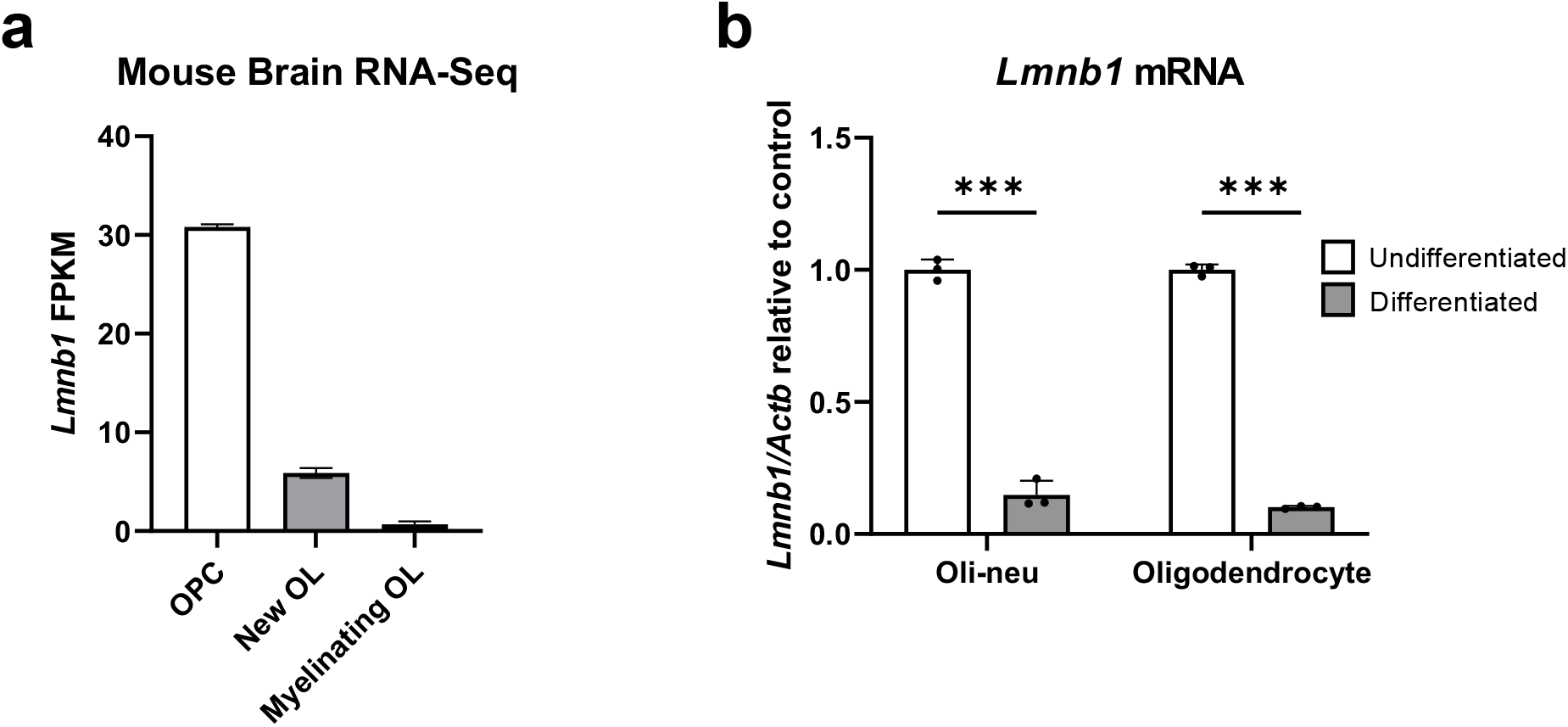
*Lmnb1* expression in oligodendrocytes is significantly reduced after differentiation. **a** Mouse Brain RNA-seq data^18^ demonstrate a significant reduction of *Lmnb1* mRNA in newly formed and myelinating oligodendrocytes (OL), compared to undifferentiated Oligodendrocyte Precursor Cells (OPCs). **b** Oli-neu and primary OLs show a similar decrease of *Lmnb1* mRNA transcript after differentiation. n = 3 biological replicates, ***: p < 0.001, using Student’s t-test.

**Supplementary Figure 5:**
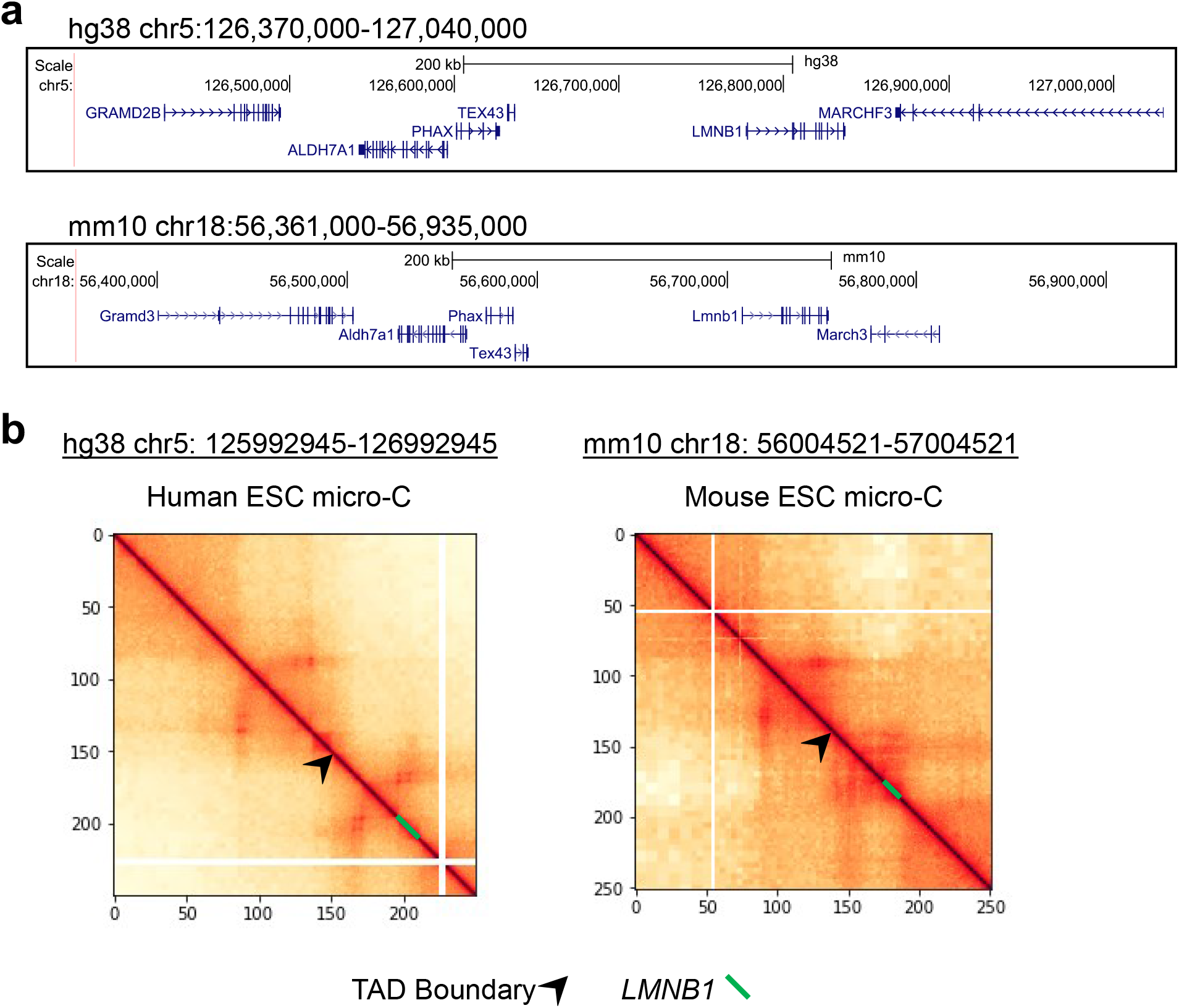
Gene order and 3D chromatin architecture surrounding the lamin B1 locus in mouse and human genomes. **a** UCSC genome browser views of the lamin B1 locus and surrounding genes in human (top) and mouse (bottom) genomes demonstrating conservation of gene order. **b** Micro-C plots showing 3D chromatin interactions and the TAD boundary (arrowhead) and the lamin B1 gene (green line) in mouse and human genomes.

**Supplementary Figure 6:**
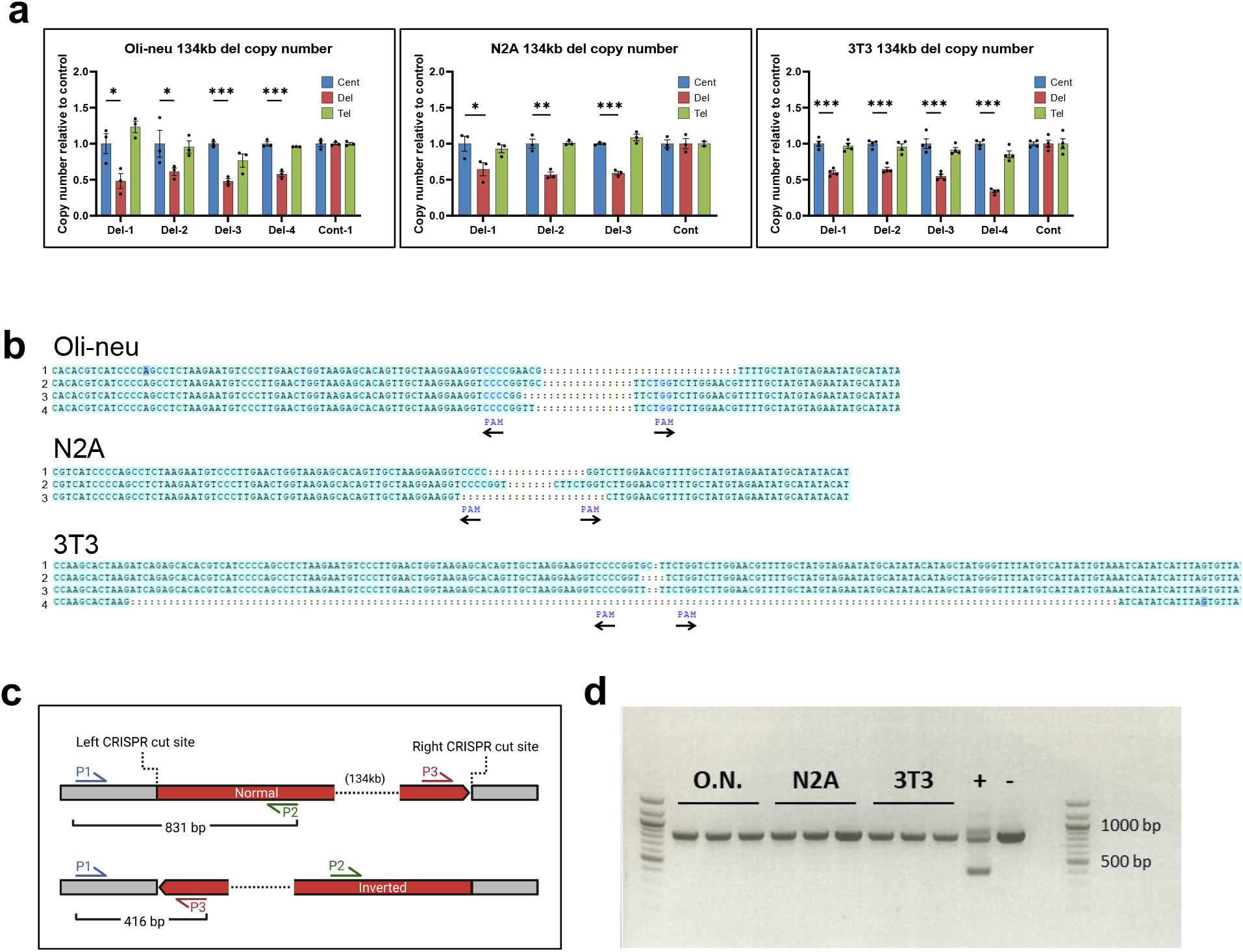
Validation of CRISPR-deleted 134 kb clones. **a** Real-time PCR genomic DNA analysis measuring relative copy number across oli-neu, N2A, and 3T3 cell lines with unique 134 kb CRISPR-mediated deletions relative to control cells. ‘Del’ measures relative copy number within the deleted region, ‘Cent’ measures relative copy number upstream of the deleted region (toward the centromere), and ‘Tel’ measures relative copy number downstream of the deleted region (toward the telomere). Graphs are mean ± SEM across n ≥ 3 technical replicates for each unique clone. *: p < 0.05, **: p < 0.01, ***: p < 0.001, calculated using Student’s t-test. **b** Aligned Sanger sequences of deletion junctions in oli-neu, N2A and 3T3 cell lines with unique 134 kb deletions. Gaps in the middle delineated by colons represent the location of deleted sequence, with PAM marking the location of the remaining spCas9 protospacer adjacent motifs ‘NGG’. Arrows denote their orientation. **c** Schematic of PCR to test for inversions. Expected band sizes are 831 bp in wild-type cells and 416 bp for cells with an inversion of the 134 kb deleted region. **d** Agarose gel of inversion PCR products, showing that there are no inversions detected in the cell lines tested. ‘+’ represents Genomic DNA from cell lines previously identified to have the 134 kb inversion. ‘-’ represents wild-type mouse genomic DNA.

**Supplementary Figure 7:**
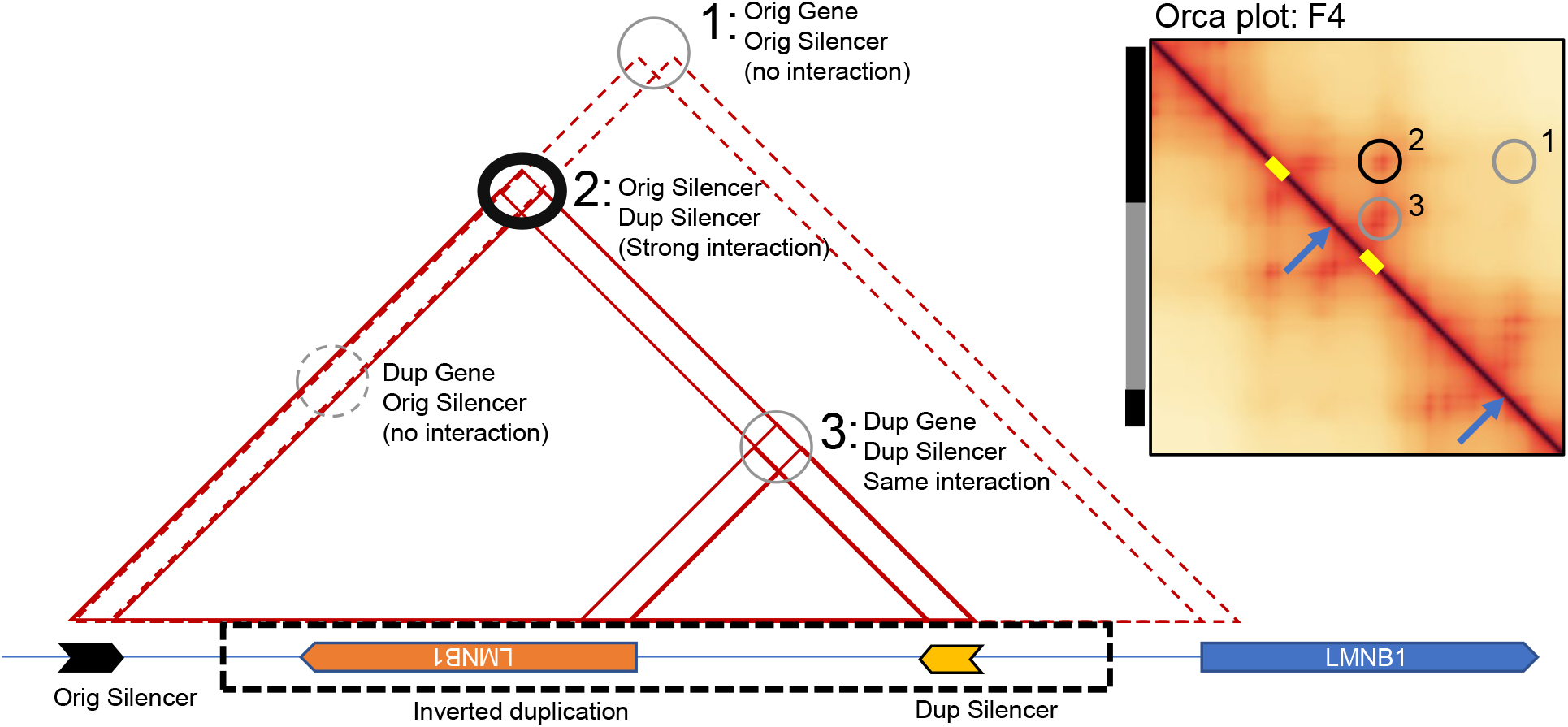
3D chromatin architecture of the ADLD inverted duplication. Model of 3D chromatin interactions in the ADLD inverted duplication reveals the formation of a novel interaction (circle 2) between the duplicated and original 19kb regulatory element (schematic is not drawn to scale). Circle 1 represents interaction between the original (non-duplicated) *LMNB1* promoter and its cognate 19 kb regulatory element and is significantly reduced. Circle 3 represents the interaction between the inverted *LMNB1* promoter and its cognate 19 kb regulatory element and is unchanged. This results in a potential loss of interaction strength between the promoter of the non-duplicated *LMNB1* and its cognate 19 kb regulatory element, as these interactions are mutually exclusive.

**Supplementary Figure 8:**
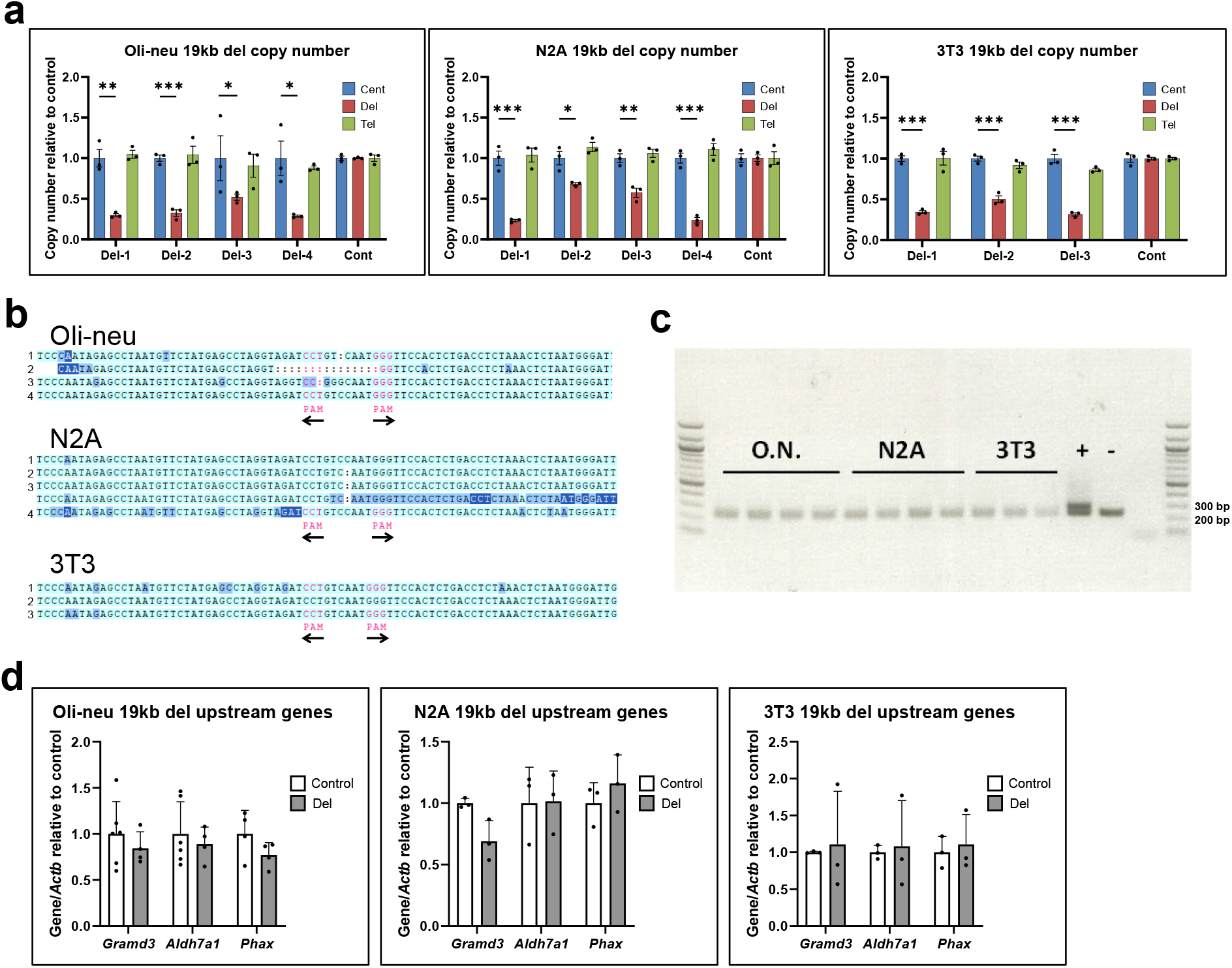
Validation of CRISPR-deleted 19 kb clones. **a** Real-time PCR genomic DNA analysis measuring relative copy number across oli-neu, N2A, and 3T3 cell lines with unique 19kb CRISPR-mediated deletions relative to control cells. ‘Del’ measures relative copy number within the deleted region, ‘Cent’ measures relative copy number upstream of the deleted region (toward the centromere), and ‘Tel’ measures relative copy number downstream of the deleted region (toward the telomere). Graphs are mean ± SEM across n = 3 technical replicates for each line. *: p < 0.05, **: p < 0.01, ***: p < 0.001, calculated using Student’s t-test. **b** Aligned Sanger sequences of deletion junctions in oli-neu, N2A, and 3T3 cell lines with unique 19kb CRISPR-mediated deletions. Gaps in the middle delineated by colons represent the location of deleted sequence, with PAM marking the location of the remaining spCas9 protospacer adjacent motifs ‘NGG’. Arrows denote their orientation. **c** Agarose gel of inversion PCR products, showing that there are no inversions detected in the cell lines tested. Expected band sizes are 241 bp in wild-type cells and 291 bp in cells with an inversion of the 19 kb deleted region. “+” represents genomic DNA from cell lines previously identified to have the 19kb inversion. “-” represents wild-type mouse genomic DNA. **d** Real-time PCR mRNA analysis measuring expression of nearby genes, *Gramd3*, *Aldh7a1*, and *Phax* in oli-neu, N2A, and 3T3 cell lines cells with the 19kb CRISPR-mediated deletion relative to control cells. Graphs are mean ± SEM across at least n = 3 independent clones for each group and cell type.

**Supplementary Figure 9:**
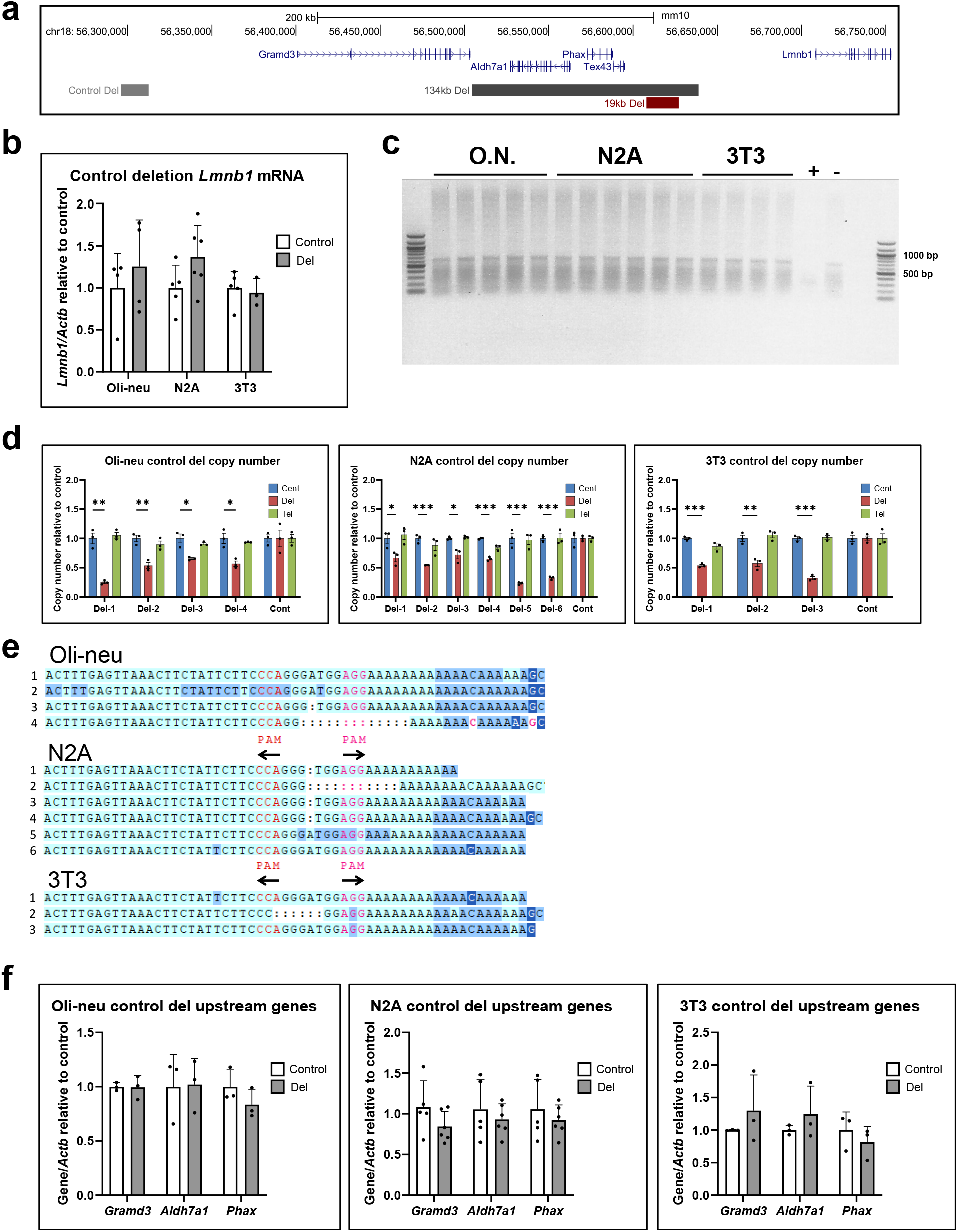
Validation of control CRISPR clones. **a** Schematic of control CRISPR deletion (grey bar) relative to the 134 kb and 19 kb deletions. The control deletion spans a 16 kb region of the genome approximately 390 kb upstream of *Lmnb1* and is outside the *Lmnb1* TAD. **b** Real-time PCR analysis of *Lmnb1* mRNA expression in oli-neu, N2A, and 3T3 cells with the 16 kb control deletion. *Lmnb1* expression is not significantly different in any cell type relative to their respective controls. *Lmnb1* expression is normalized to b actin (*Actb*), n ≥ 3 unique cell lines for both control and deletion clones for each cell type. **c** Agarose gel of inversion PCR products, showing that there are no inversions detected in the cell lines tested. Expected band sizes are 628 bp in wild-type cells and 256 bp in cells with an inversion of the 16 kb control region. “+” represents genomic DNA from cell lines previously identified to have the 16kb inversion. “-” represents wild-type mouse genomic DNA. **d** Real-time PCR genomic DNA analysis measuring relative copy number across oli-neu, N2A, and 3T3 cell lines with unique 16kb CRISPR-mediated deletions relative to control cells. ‘Del’ measures relative copy number within the deleted region, ‘Cent’ measures relative copy number upstream of the deleted region (toward the centromere), and ‘Tel’ measures relative copy number downstream of the deleted region (toward the telomere). Graphs are mean ± SEM across n = 3 technical replicates for each line. *: p < 0.05, **: p < 0.01, ***: p < 0.001, calculated using Student’s t-test. **e** Aligned Sanger sequences of deletion junctions in Oli-neu, Neuro2a, and NIH-3T3 cell lines with unique 16kb CRISPR-mediated deletions. Gaps in the middle delineated by colons represent the location of deleted sequence, with PAM marking the location of the remaining spCas9 protospacer adjacent motifs ‘NGG’. Arrows denote their orientation. **f** Real-time PCR mRNA analysis measuring *Gramd3*, *Aldh7a1*, and *Phax* expression in oli-neu, N2A, and 3T3 cells with the 16kb CRISPR-mediated deletion relative to control cells. Graphs are mean ± SEM across at least n = 3 independent clones for each group and cell type.

**Supplementary Figure 10:**
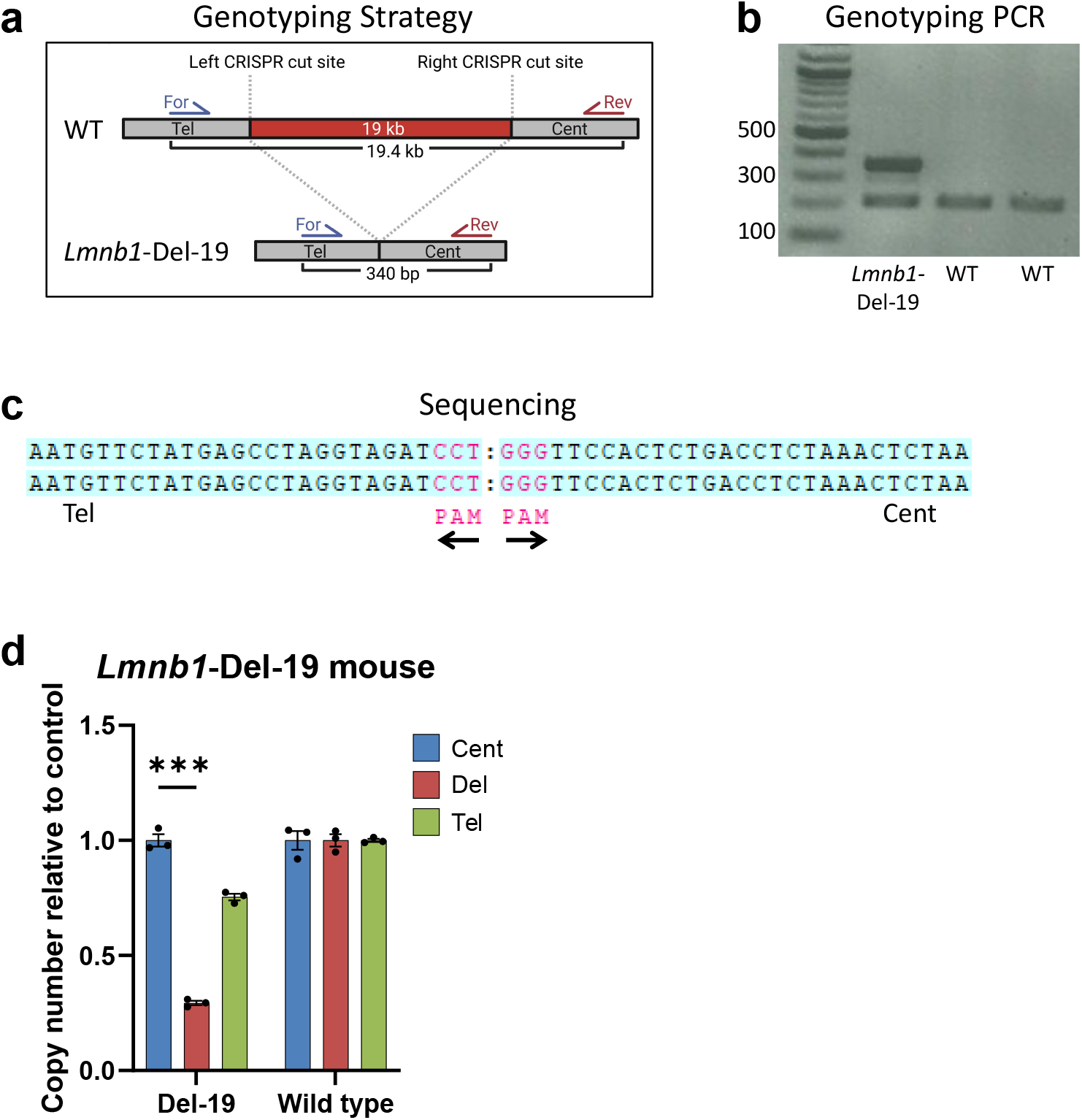
Characterization of *Lmnb1*-Del-19 mice. **a** Gentoyping strategy for *Lmnb1*-Del-19 mice. The forward and reverse primers are normally 19.4 kb apart in wild type (WT) mice but yield a 340 bp PCR product in *Lmnb1*-Del-19 mice. **b** Representative genotyping PCR of a *Lmnb1*-Del-19 mouse and two wild-type mice as controls. The band at 200 bp is an endogenous control. **c** Sequencing across the deletion junction in *Lmnb1*-Del-19 mice reveals the deletion of ∼19kb between the two PAM sites. Arrows underneath the PAM sites indicate the orientation of the protospacer and PAM sequences. **d** Copy number analysis of DNA from a *Lmnb1*-Del-19 mouse using real time PCR demonstrates reduced copy number using primers within deleted region (red). n = 3 technical replicates, *** p < 0.001. Primers outside deleted region (blue and green) show copy numbers similar to wild-type mice.

**Supplementary Figure 11:**
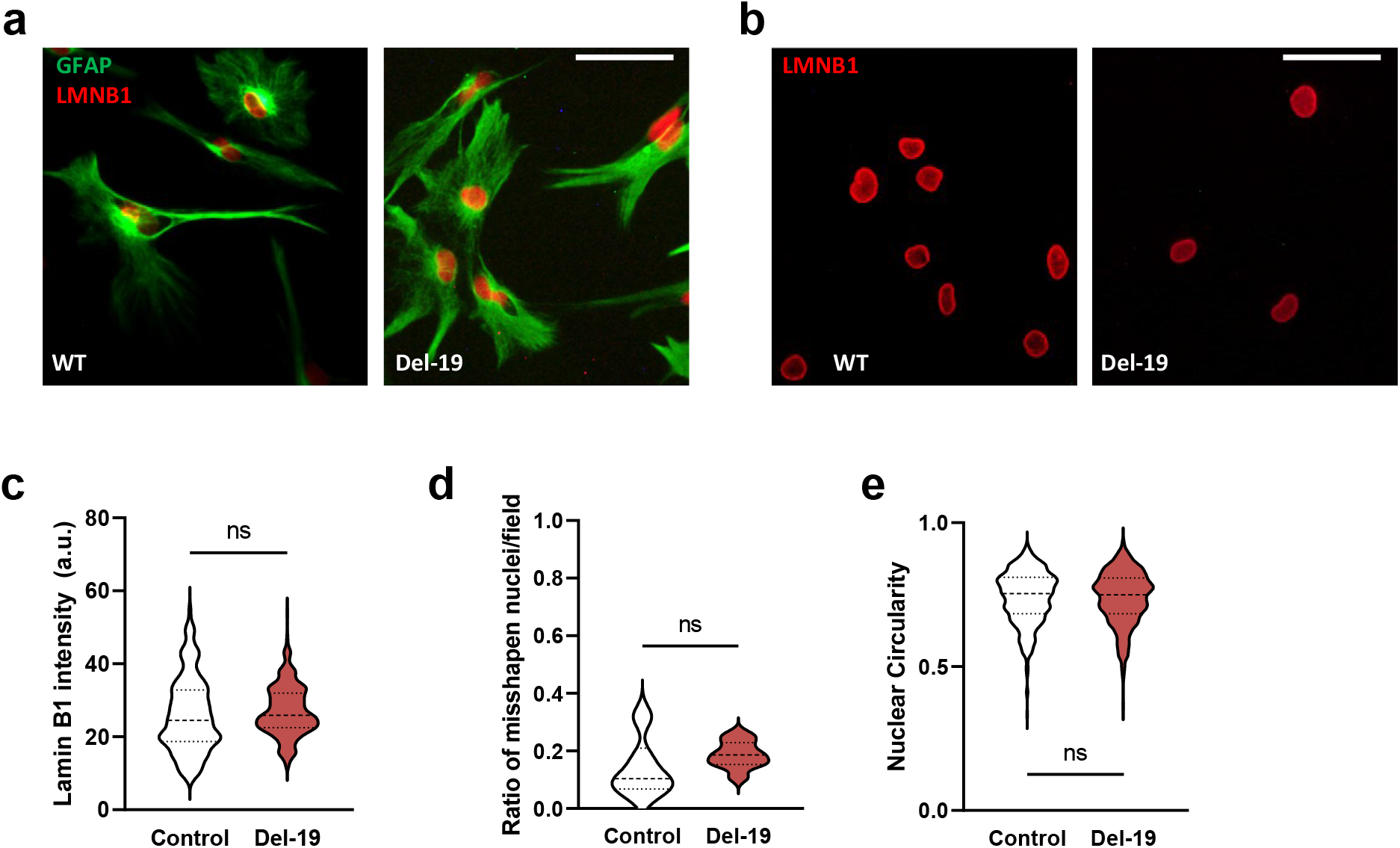
Nuclear morphology in *Lmnb1*-Del-19 mouse astrocytes. **a** Representative epifluorescence images of cultured primary differentiated astrocytes (AS) from *Lmnb1*-Del-19 (Del-19) mice and WT controls stained with antibodies against LMNB1 (red) and the astrocyte specific marker GFAP (green). Scale bar = 50*μ*m. **b** Staining of OL nuclei from Del-19 mice and WT demonstrate no differences of LMNB1 intensity or nuclear shape between WT and Del-19 AS nuclei. Scale bar = 50*μ*m. Violin plot quantification of **c** LMNB1 intensity, **d** ratio of misshapen nuclei, and **e** circularity reveal no significant differences between astrocyte nuclei from Del-19 mice relative to WT, calculated using two-tailed t-test (b) and two-tailed Mann-Whitney test (c and d).

**Supplementary Figure 12:**
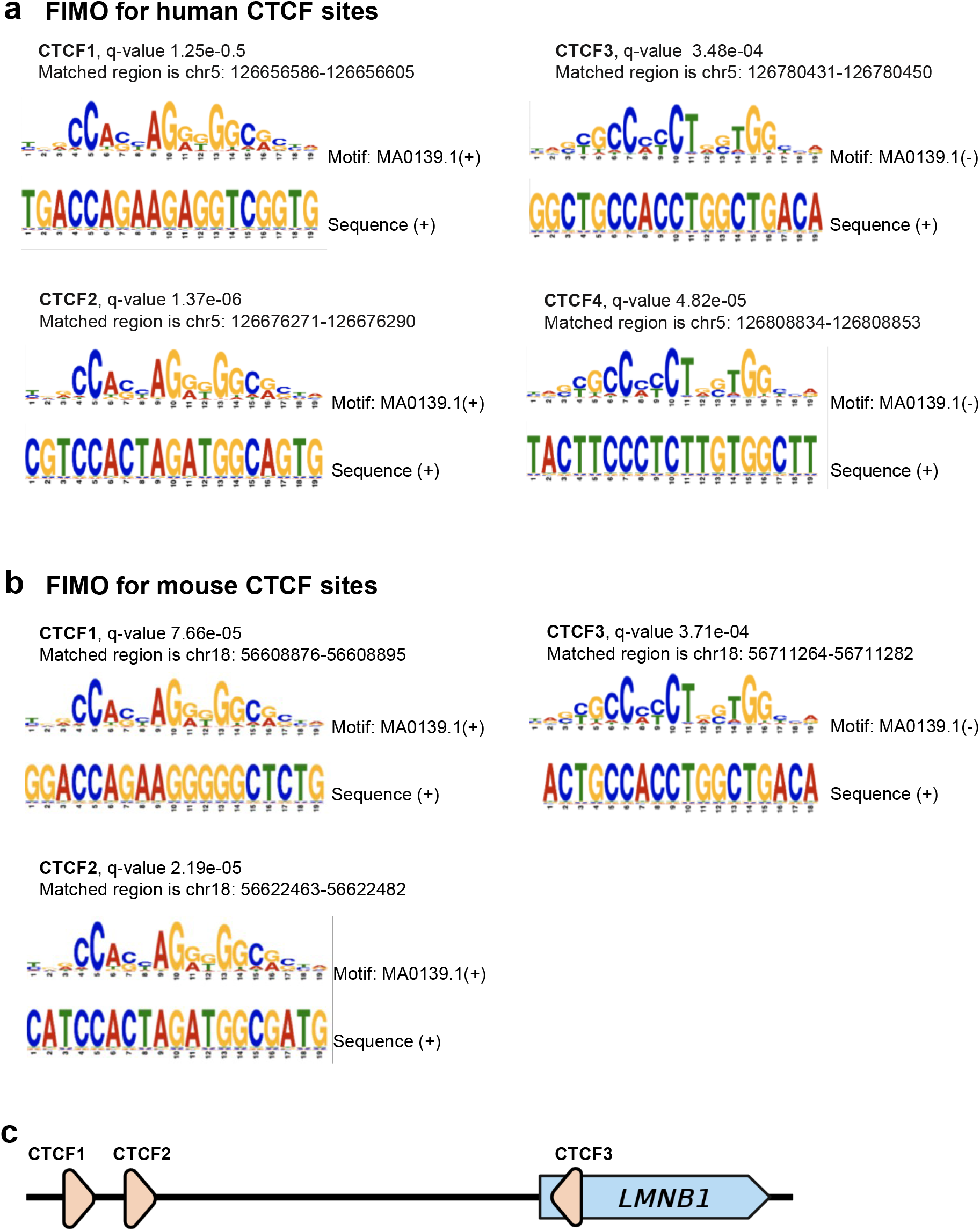
Analysis of CTCF motifs. Find Individual Motif Occurrences (FIMO) analysis was applied search for CTCF binding sites within the critical region for **a** human and **b** mouse genomes. The discovered matches and p-values are shown. “Normal” and “Reverse Complement” indicate sequence directionality. **c** graphical representation of CTCF1, 2, and 3 sites and their orientation with respect to Lamin B1 (applies to both mouse and human, not to scale).

**Supplementary Figure 13:**
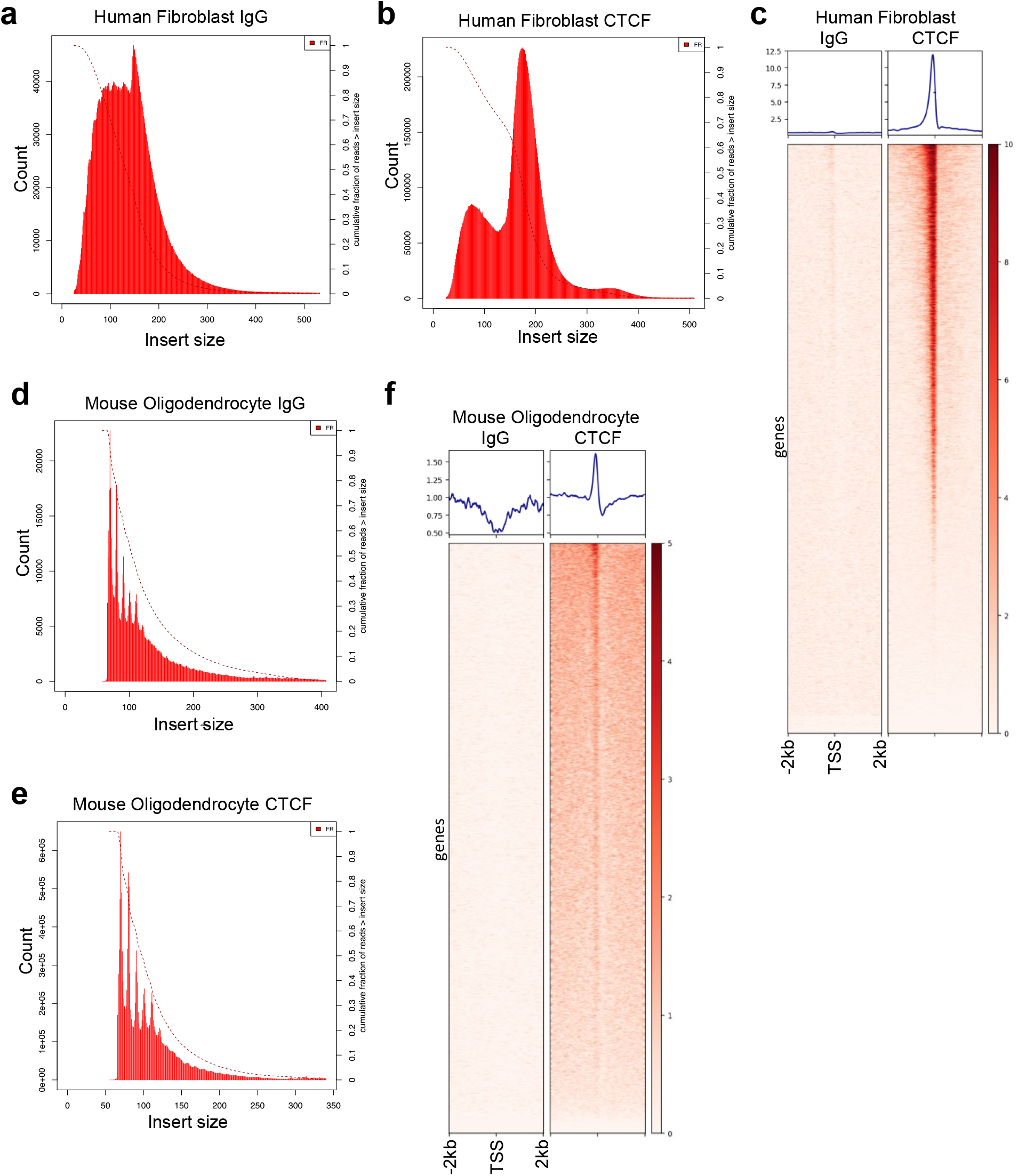
CTCF CUT&RUN quality controls. **a-b, d-e** Insert size plots showing read distribution among combined replicates for each condition and sample type. Insert sizes were calculated using Picard after removing duplicate and low-quality reads but before performing any read normalization. Dashed lines indicate fraction of reads greater than insert size (right y-axis). **c** Human fibroblast CTCF CUT&RUN visualized over all hg38 transcription start sites. **f** Mouse Oligodendrocyte CTCF CUT&RUN visualized over all mm10 transcription start sites. IgG serves as a negative control experiment to assess background cutting.

**Supplementary Figure 14:**
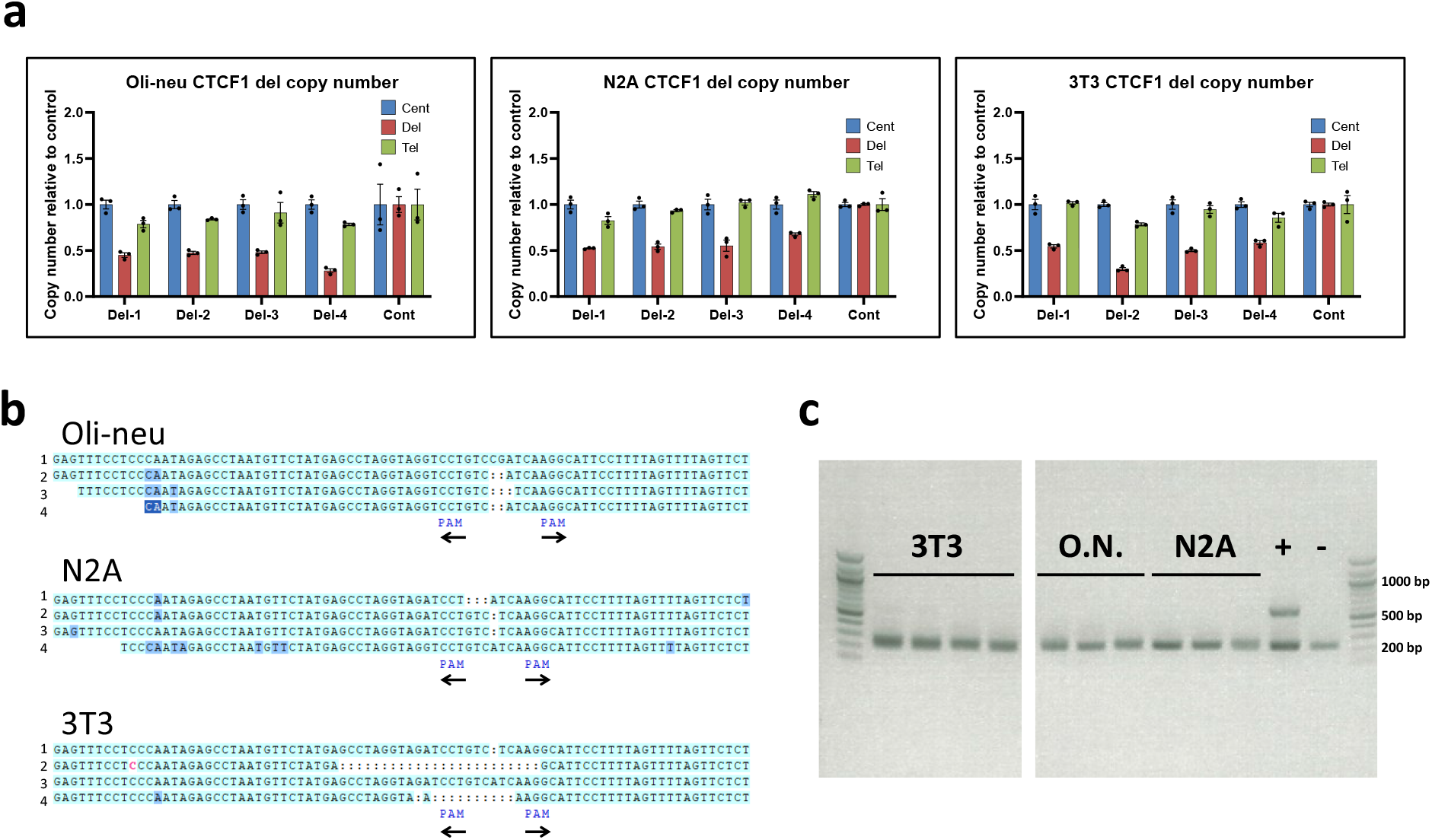
Validation of CRISPR deleted CTCF1 clones. **a** Real-time PCR genomic DNA analysis measuring relative copy number across oli-neu, N2A and 3T3 cell lines with unique CTCF1 CRISPR-mediated deletions relative to control cells. ‘Del’ measures relative copy number within the deleted region, ‘Cent’ measures relative copy number upstream of the deleted region (toward the centromere), and ‘Tel’ measures relative copy number downstream of the deleted region (toward the telomere). Graphs are mean ± SEM across n = 3 technical replicates for each line. *: p < 0.05, **: p < 0.01, ***: p < 0.001, calculated using Student’s t-test. **b** Aligned Sanger sequences of deletion junctions in oli-neu, N2A and 3T3 cell lines with unique CTCF1 CRISPR-mediated deletions. Gaps in the middle delineated by colons represent the location of deleted sequence, with PAM marking the location of the remaining spCas9 protospacer adjacent motifs ‘NGG’. Arrows denote their orientation. **c** Agarose gel of inversion PCR products, showing that there are no inversions detected in the cell lines tested. Expected band sizes are 240 bp in wild-type cells and 596 bp in cells with an inversion of the CTCF1 deleted region. “+” represents genomic DNA from cell lines previously identified to have the inversion. “-” represents wild-type mouse genomic DNA.

**Supplementary Fig. 15:**
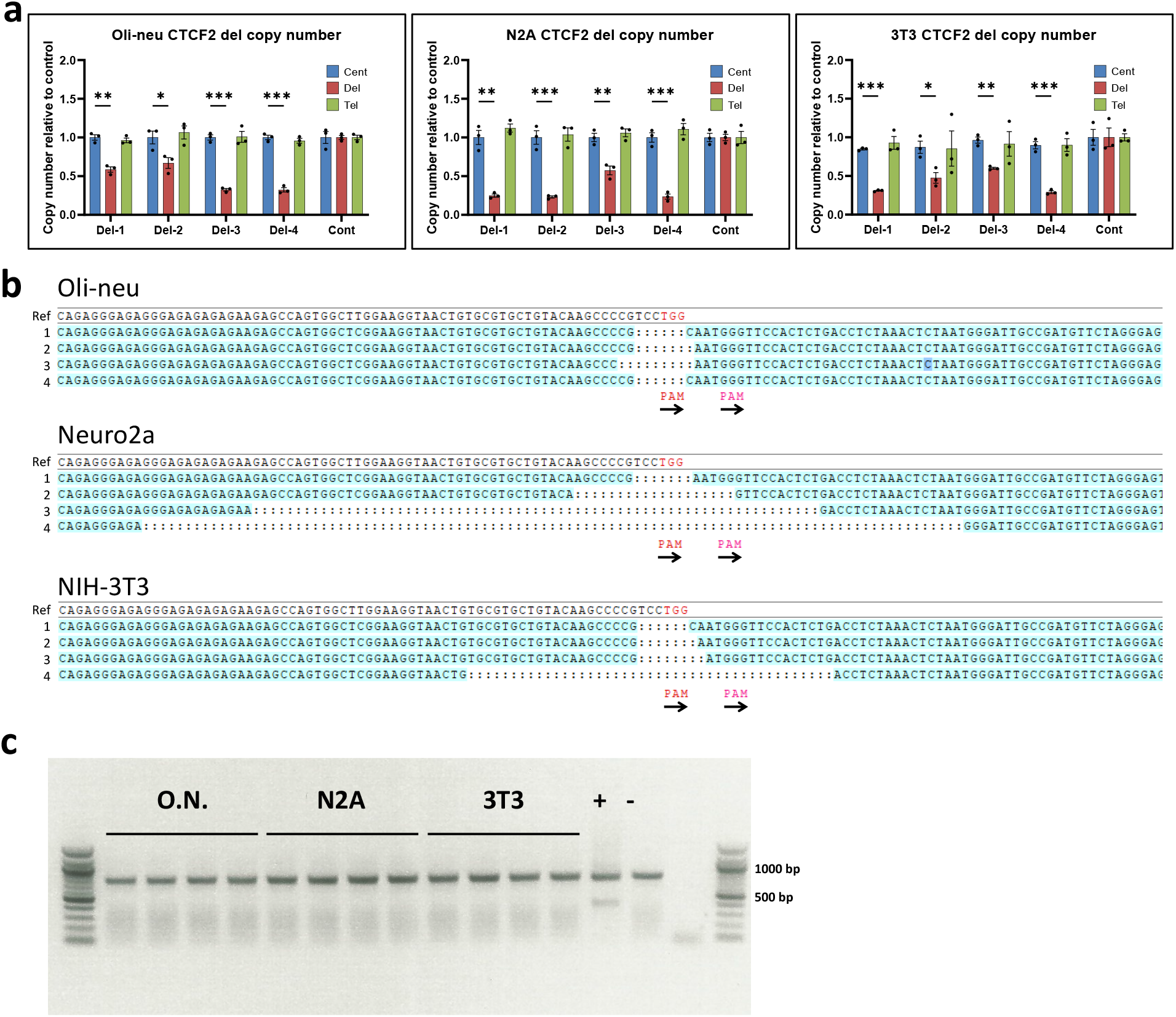
Validation of CRISPR deleted CTCF2 clones. **a** Real-time PCR genomic DNA analysis measuring relative copy number across oli-neu, N2A and 3T3 cell lines with unique CTCF2 CRISPR-mediated deletions relative to control cells. ‘Del’ measures relative copy number within the deleted region, ‘Cent’ measures relative copy number upstream of the deleted region (toward the centromere), and ‘Tel’ measures relative copy number downstream of the deleted region (toward the telomere). Graphs are mean ± SEM across n = 3 technical replicates for each line. *: p < 0.05, **: p < 0.01, ***: p < 0.001, calculated using Student’s t-test. **b** Sanger sequences of deletion junctions in oli-neu, N2A and 3T3 cell lines with unique CTCF2 CRISPR-mediated deletions. Sequences have been aligned to a reference sequence (Ref, top) Gaps in the middle delineated by colons represent the location of deleted sequence, with PAM marking the location of the remaining spCas9 protospacer adjacent motifs ‘NGG’. Arrows denote their orientation. **c** Agarose gel of inversion PCR products, showing that there are no inversions detected in the cell lines tested. Expected band sizes are 768 bp in wild-type cells and 462 bp in cells with an inversion of the CTCF2 deleted region. “+” represents genomic DNA from cell lines previously identified to have the inversion. “-” represents wild-type mouse genomic DNA.

**Supplementary Figure 16:**
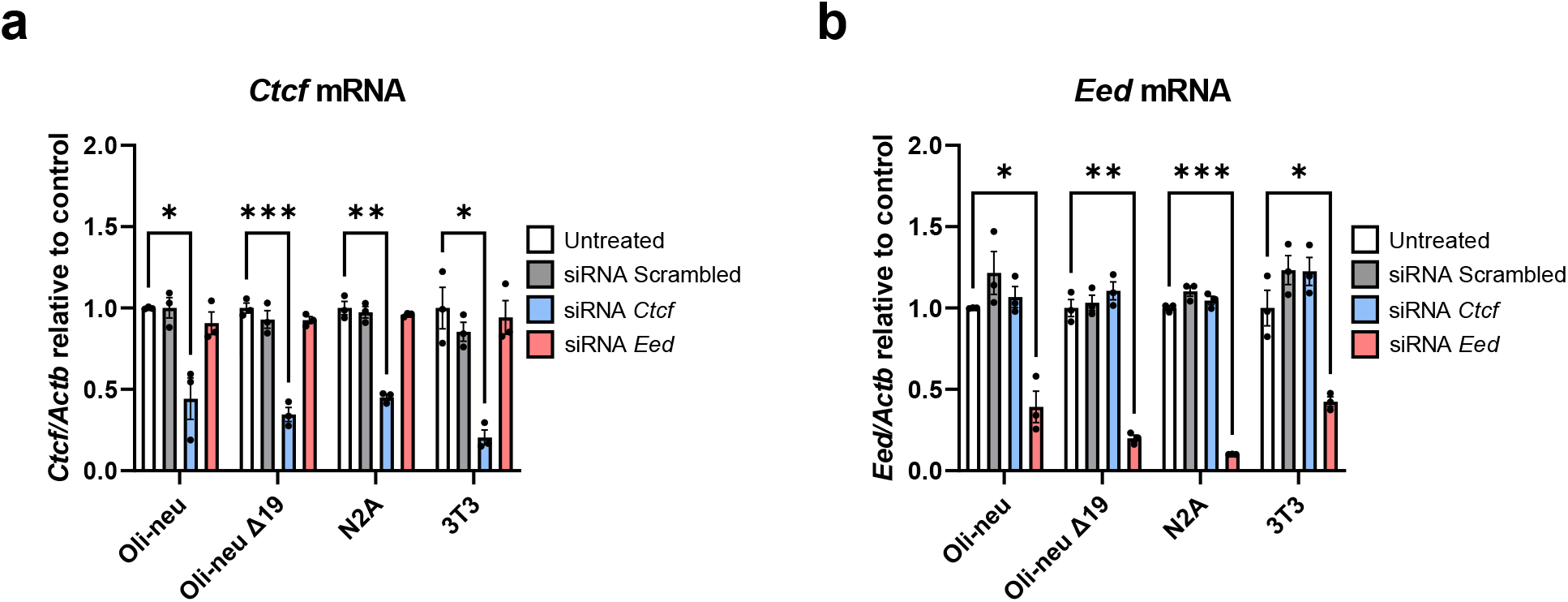
*Ctcf* and *Eed* siRNA treatment. Real-time PCR on **a** *Ctcf* and **b** *Eed* mRNA from wild-type oli-neu, N2A, 3T3, and Δ19 oli-neu cells treated with scrambled siRNA and siRNA against *Ctcf* and *Eed*, demonstrate siRNA-mediated knockdown is specific for their respective genes. Graphs are mean ± SEM for 3 biological replicates per cell type and treatment. *: p < 0.05, **: p < 0.01, ***: p < 0.001, using two-way ANOVA.

**Supplementary Figure 17:**
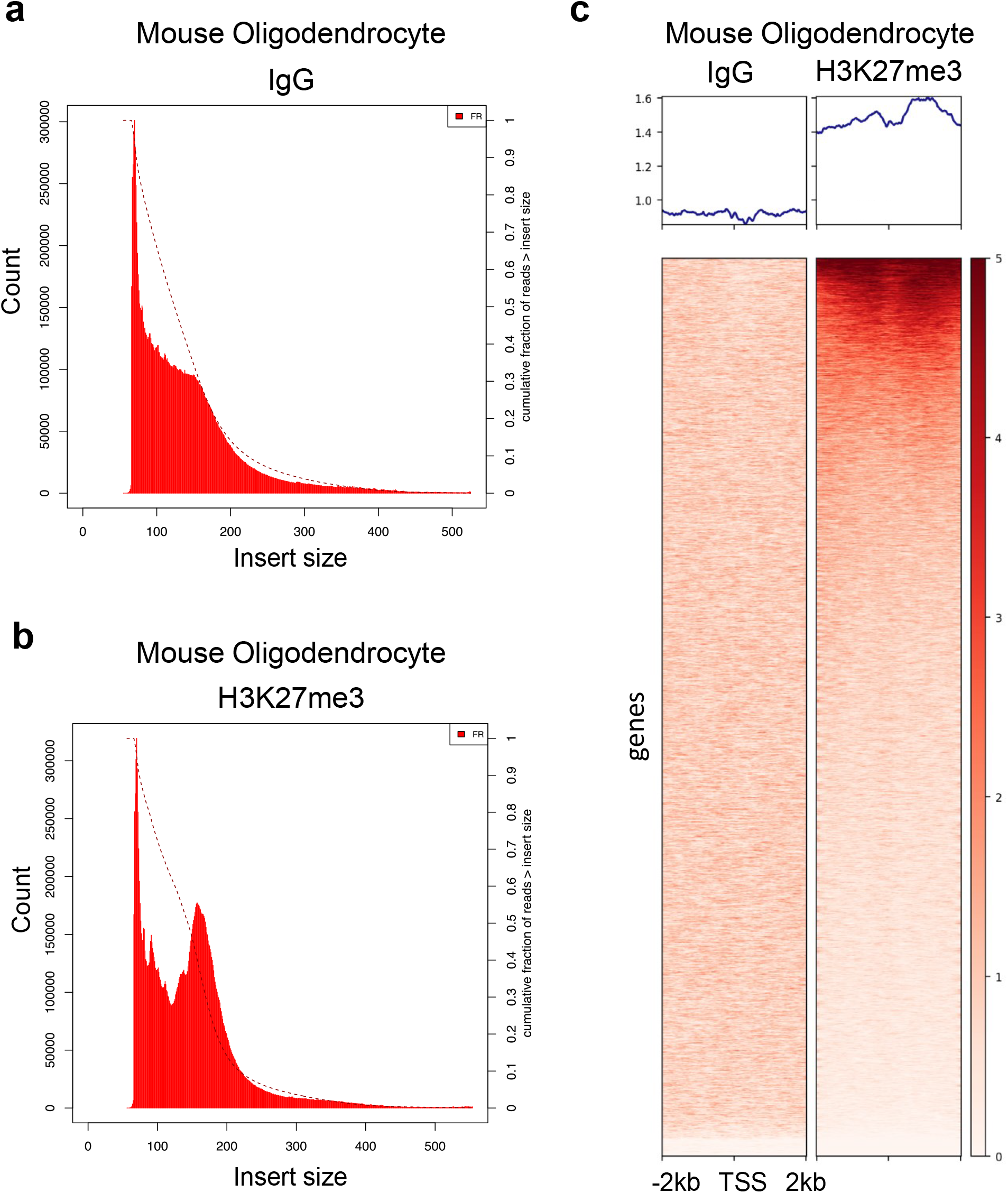
H3K27me3 CUT&RUN quality controls. **a-b** Insert size plots showing read distribution among combined replicates for each condition. Insert sizes were calculated using Picard after removing duplicate and low-quality reads but before performing any read normalization. Dashed lines indicate fraction of reads greater than insert size (right y-axis). **c** Mouse Oligodendrocyte H3K27me3 CUT&RUN visualized over all mm10 transcription start sites. IgG serves as a negative control experiment to assess background cutting.

**Supplementary Figure 18:**
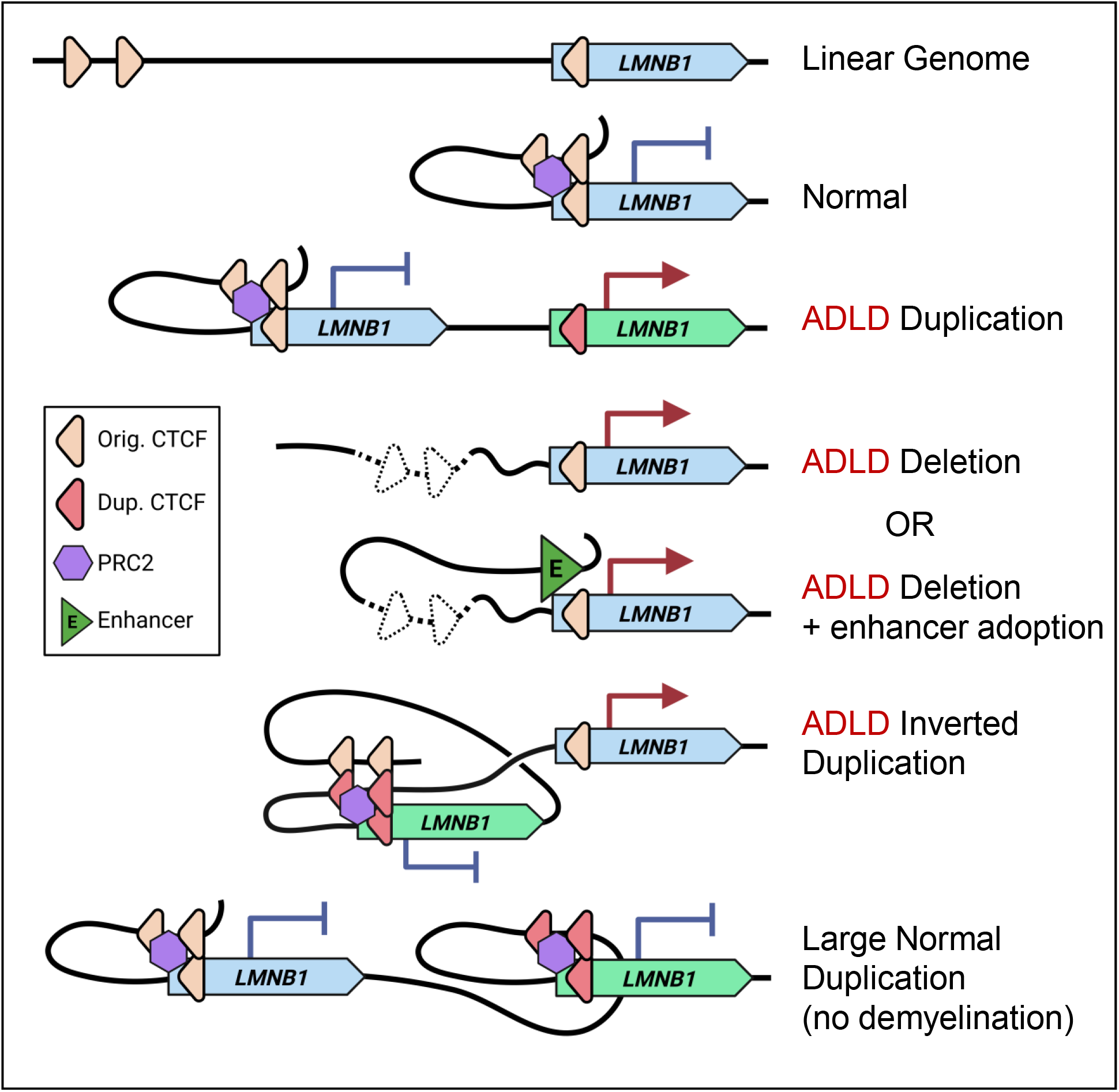
Model of OL-specific silencing of *LMNB1*. Oligodendrocyte-specific silencing of LMNB1 is mediated by CTCF binding and PRC2 complex recruitment. Normally, *LMNB1* is silenced when upstream CTCF sites interact with the CTCF site in the *LMNB1* gene, recruiting PRC2 complex proteins and attenuating transcription. In the case of ADLD-Dup cases, the duplicated *LMNB1* copy lacks CTCF interactions, losing transcriptional silencing and leading to OL specific overexpression and disease. A loss of the upstream CTCF region also results in a loss of transcriptional silencing in ADLD-Del cases. Whether the interaction of the ectopic enhancer with the *LMNB1* promoter is also required for disease causation is unclear and both possibilities are presented. LN-Dups cases duplicate both the upstream CTCF sites in addition to the *LMNB1* gene, thereby preserving the interactions that lead to transcriptional silencing. In the case of inverted duplications, our results predict that the duplicated *LMNB1* copy is silenced by its cognate duplicated CTCF binding sites. However, its insertion disrupts the interaction between the original CTCF sites and non-duplicated *LMNB1* gene. This would result in OL specific overexpression as with canonical the ADLD-Dup and Del configurations. The original CTCF site also forms a novel interaction with the duplicated CTCF site further reducing the availability of the former with the non-duplicated gene leading to an even greater disruption of transcriptional silencing and potentially a more severe disease phenotype.

