## Supplementary tables for "An oligodendrocyte silencer element underlies the pathogenic impact of lamin B1 structural variants"

#### Supplementary Table 1: Details of *LMNB1* duplications

| Family | Duplication coordinates (hg38) | Dup Size (bp) | Microhomology |
|---|---|---|---|
| F1 | chr5:122477254-126843256 | 4366003 | 49 bp homology |
| F2 | chr5:125918128-126949480 | 1031353 | C-insertion |
| F3 | chr5:126512007-126978629 | 466623 | AG |
| F4 | chr5:126363827-126838825 | 474999 | Complex inversion |

### Supplementary Table 2: Oligonucleotides used for the study

| | **Array CGH Junction PCR** | |
|---|---|---|
| | Primer Sequences | Expected Size |
| Family F1 | Forward (telomeric end): AATCAGGCAGTGTGAGTCTTCA | 1.45 kb |
| | Reverse (centromeric end): CCCCAATCGCTTATTTCTGCAC | |
| Family F2 | Forward (telomeric end): CTTTGTCGGCCTACAGTGGT | 9.06 kb |
| | Reverse (centromeric end): TCCCTTCCAGGGTCTCTTTTA | |
| Family F3 | Forward (telomeric end): CAACACTATTCGGCCATTCC | 1.51 kb |
| | Reverse (centromeric end): GTCAGTCAGAACCACAGGTCAC | |

| | **CRISPR gRNA design** | |
|---|---|---|
| | Spacer Sequences | PAM |
| 134kb deletion: | GAGATGGCAAAATCGCACCG | GGG |
| | TTAGAATCCGCTGTCACTTC | TGG |
| 19kb deletion: | GGATTGCACAATCACCGGAC | AGG |
| | GGAGTTGTAACGGCGGCAAT | GGG |
| CTCF1 deletion: | GGATTGCACAATCACCGGAC | AGG |
| | ATTGATTGGTGACATCATCA | AGG |
| CTCF2 deletion: | GTGCTGTACAAGCCCCGTCC | TGG |
| | GGAGTTGTAACGGCGGCAAT | GGG |
| Control deletion: | CCATTGTACAACCCACCCCT | GGG |
| | TATTCACCTATGTCCAATGG | AGG |

| | **Screening deletion junctions and sequencing** | |
|---|---|---|
| | Primer Sequences | Expected Size |
| 134kb deletion: | GGTGGAGCCTCTAAAGCATGTA | 670bp |
| | TTTAGTCTCGGCTGCTTCTCTC | |
| 19kb deletion: | AAACCACCACAACTCCTTTCAG | 349bp |
| | CAGCTAAGGTGGGGGTCTTA | |
| CTCF1 deletion: | AAACCACCACAACTCCTTTCAG | 240bp |
| | ACCAACAGAACACAAACAGCAC | |
| CTCF2 deletion: | CGAGTGACTTAGGGACCTTGAC | 1866bp |
| | CAGCTAAGGTGGGGGTCTTA | |
| Control deletion: | GTGTAGAAAGGAGGCTAGCGAT | 892bp |
| | GCAGGAGGTTTGGGTATCAGTT | |

### Supplementary Table 2 (continued)

| | **Testing for genomic inversions** | |
|---|---|---|
| | Primer Sequences | Expected Size |
| 134kb: | P1: GGTGGAGCCTCTAAAGCATGTA | Normal: 831bp<br>Inverted: 416bp |
| | P2: GGTTTTCGGTCTTTCTTCCCAC | |
| | P3: GTGACTTAGACCCAGACATGCA | |
| 19kb: | P1: AAACCACCACAACTCCTTTCAG | Normal: 241bp<br>Inverted: 291bp |
| | P2: TCGAGGTTCCTCAGACAGACA | |
| | P3: CTCCTTTCCCTGTCTTCTTCCT | |
| CTCF1: | P1: AAACCACCACAACTCCTTTCAG | Normal: 240bp<br>Inverted: 596bp |
| | P2: TCGAGGTTCCTCAGACAGACA | |
| | P3: TGTTGTGGTGCTCTTTTCCTGA | |
| CTCF2: | P1: CTGACTATGGCTTAAGGGGAAA | Normal: 768bp<br>Inverted: 462bp |
| | P2: GTAGACTTGATTCTCGCCTTCC | |
| | P3: CTCCTTTCCCTGTCTTCTTCCT | |
| Control: | P1: GAATAATGATGGTGAAGGCT | Normal: 628bp<br>Inverted: 256bp |
| | P2: ATTAAGAGACTTATAGTTCAGT | |
| | P3: ACACTAAGTTCTTCTGCCCAGG | |

| | **Generation of Lmnb1-del-19 mice** |
|---|---|
| | Oligonucleotide Sequences |
| Left sgRNA For | TGTAATACGACTCACTATAGGGGATTGCACAATCACCGGACGTTTTAGAGCTAGAAATAGC |
| Right sgRNA For | TGTAATACGACTCACTATAGGGGAGTTGTAACGGCGGCAATGTTTTAGAGCTAGAAATAGC |
| Universal Rev | AAAAGCACCGACTCGGTGCC |

| | **mRNA expression qPCR primers** |
|---|---|
| | Primer Sequences |
| Human *ACTB*: | GGACCTGACTGACTACCTCAT |
| | CGTAGCACAGCTTCTCCTTAAT |
| Human *LMNB1*: | GATTGCCCAGTTGGAAGCCT |
| | TGGTCTCGTTAATCTCCTCTTCATACA |
| Mouse *Actb*: | CCACTGCCGCATCCTCTTCC |
| | CTCGTTGCCAATAGTGATGACCTG |
| Mouse *Lmnb1*: | GCTGCTCAATTATGCCAAGAAG |
| | GCCGCATCCTTAGAGTTTAGT |
| Mouse *Gramd3*: | CTGCTGCTGCCTGAAGATAA |
| | GGAGTGCTGTGTTTCCTGAT |
| Mouse *Aldh7a1*: | GGATAACGAGGGCGTGTATAA |
| | CCTGTCGGACTCTTGCTATT |
| Mouse *Phax*: | ATAGCCCGAGTAGTGAGGATAC |
| | TGTTCTTCTTCGGCTACCATTC |

### Supplementary Table 2 (continued)

| | **Copy number analysis qPCR primers** |
|---|---|
| | Primer Sequences |
| Cent (134kb/ 19kb/ CTCF1/ CTCF2): | CTGCTGCTGCCTGAAGATAA |
| | GGAGTGCTGTGTTTCCTGAT |
| Cent (Control): | TGAGAGCAACTTGGGAGAAAG |
| | AGTACGCTTGGAGTCCTAGAT |
| Del (134kb/ 19kb/ CTCF1): | CTCACCTTTCCTTCCTGACATT |
| | CCACGCCTGTTTCCTCTTTA |
| Del (CTCF2): | TGGAGCATTACAGTGTGACTAAT |
| | CGCTGACCTTGAGAGTCTTATC |
| Del (Control): | GGGCAAGATTAGAGATGGTTAGAG |
| | GTAGGTCTTCCAACACACTTCA |
| Tel (all): | GTCAGGAGAGAACTGGAAAGAC |
| | GTTCCCATTGAGAAGGGAGATT |

**Supplementary Table 3: Antibodies used for the study**

| Primary antibodies | Species | Application | Manufacturer | Catalog # |
|---|---|---|---|---|
| CNPase | Mouse | WB | Millipore | NE1020 |
| FLAG | Mouse | WB | Sigma | F1804 |
| GAPDH | Mouse | WB | ThermoFisher | MA5-15738 |
| GFAP | Mouse | ICC/IF | Millipore | MAB360 |
| Lamin B1 | Rabbit | WB, ICC/IF | Abcam | ab16048 |
| PDGFRα | Rat | Immunopan | BD Pharmingen | 562171 |
| K27me3 | Rabbit | CUT&RUN | Millipore | 07-449 |
| CTCF | Rabbit | CUT&RUN | Millipore | 07-729 |

| Secondary antibodies | Species | Application | Manufacturer | Catalog # |
|---|---|---|---|---|
| Anti-Mouse AlexaFluor-488 | Donkey | ICC/IF | Jackson Immunoresearch | 715-545-150 |
| Anti-Mouse Cy3 | Donkey | ICC/IF | Jackson Immunoresearch | 715-165-150 |
| Anti-Mouse IRDye 680LT | Goat | WB | LI-COR Biosciences | 926-68020 |
| Anti-Rabbit AlexaFluor-488 | Donkey | ICC/IF | Jackson Immunoresearch | 711-545-152 |
| Anti-Rabbit AlexaFluor-647 | Donkey | ICC/IF | Jackson Immunoresearch | 711-605-152 |
| Anti-Rabbit IRDye 800CW | Donkey | WB | LI-COR Biosciences | 926-32213 |
| Anti-Rat | Goat | Immunopan | Jackson Immunoresearch | 112-005-167 |
